# A Single-Cell and Spatial 3D Multi-omic Atlas of Developing Human Basal Ganglia and Inhibitory Neurons

**DOI:** 10.64898/2026.01.28.702385

**Authors:** Matthew G. Heffel, Heng Xu, Oier Pastor-Alonso, Xinzhe Li, Mohammad S. Baig, Rayyan Irfan Ghoor, Runjia Li, Colin Kern, Joonho Kum, Yi Zhang, Jon Paino, Min Jen Tsai, Chu-Yi Tai, Grant Tucker, Zoey Zhao, Angie Hou, Zachary von Behren, Mohini Bhade, Siqian Li, Kadellyn Sandoval, Jessica Scholes, Felicia Codrea, Jeffrey Calimlim, Emma K. Liao, Gwyneth Leung, JaeYeon Kim, Eleazar Eskin, Jonathan Flint, Jennifer A. Cotter, Bogdan Pasaniuc, Bogdan Bintu, Quan Zhu, Eran A. Mukamel, Jason Ernst, Mercedes F. Paredes, Chongyuan Luo

## Abstract

The human basal ganglia (BG), subcortical nuclei fundamental to motor regulation and cognitive modulation, is constructed from neurons produced during gestation in the adjacent ganglionic eminences (GEs). GEs are transient structures in the ventral prenatal brain that also generate GABAergic inhibitory neurons which migrate to destinations in the BG, cortex and other destinations. This study aims to elucidate the epigenomic and 3D-genomic dynamics involved in the specification and maturation of GEs and GE-derived neurons, using single-nucleus methyl-3C sequencing (snm3C-seq), highly-multiplexed spatial transcriptomics, and chromatin+RNA single-molecule imaging. Our multi-modal data support a heterogeneous temporal progression across GE subregions, with the lateral GE (LGE) showing declining neurogenic activity in mid-gestation and caudal GE (CGE) exhibiting ongoing developmental progression through infancy. We identified regulatory programs that specify subtypes of BG principal cells, medium spiny neurons (MSN), via synchronized maturation of the 3D-epigenome. In infant brains, we found a transient short-range enriched (SE) chromatin conformation during the transition between oligodendrocyte progenitors (OPCs) and oligodendrocytes (ODCs), and a temporary shift toward Long-range Enriched (LE) chromatin conformation in projection neurons, extending previous works showing the differentiation of neurons and glial cells is associated with permanent SE and LE conformation, respectively. Lastly, we found that gene regulatory regions active in MSNs were enriched in loci associated with genetic risk for neuropsychiatric disease. Our study delineates the highly complex, lineage-specific 3D genomic dynamics in ventral progenitors and basal ganglia populations of the perinatal human brain.

**Highlights:**

- Joint 3D genome and DNA methylome analysis of ventral brain progenitor zones
- Heterogeneous developmental progressions of the ganglionic eminences
- Distinct development dynamics and regulatory landscape of MSNs and interneurons
- Transient remodeling of the 3D-genome in neurons and oligodendrocyte progenitors

## Introduction

Human brain development requires progenitor cells at ventricular regions to generate distinct cellular types that migrate to the laminated cortical structures and subcortical nuclei^1^. Great advances have been made in defining the formation of cortical areas from progenitors in the dorsal ventricular zone, which primarily contribute to the excitatory lineage^1^. However, questions remain regarding the regulation of proliferation and cell fates in ventral neurogenic regions, collectively termed the ganglionic eminences (GEs)^2–4^. These embryonic territories generate inhibitory populations for both cortical and subcortical destinations. A critical target for GE-derived populations is the basal ganglia (BG), which includes the striatum and the globus pallidus. The BG is a central component of recurrent cortico–basal ganglia–thalamo–cortical circuits that support associative, limbic, and sensorimotor processing, regulating motor control and modulating mood and executive functions^5^. Yet, the progenitor patterning, developmental timing, and gene regulatory dynamics that generate the diverse inhibitory neurons within these interconnected circuits remain poorly understood.

The GEs give rise to inhibitory neurons that release Gamma-aminobutyric acid (GABA), including GABAergic projection neurons in the BG and locally projecting interneurons in the cortex and olfactory bulb. The lateral ganglionic eminence (LGE) produces medium spiny neurons (MSNs)^6^, GABAergic projection neurons that account for the vast majority of neurons in the striatum, as well as olfactory bulb interneurons^7^ and amygdala intercalated cells^8^. The MGE and CGE produce GABAergic interneurons that migrate to the cortex, hippocampus, striatum, and amygdala^9^. The migration of MGE and CGE-derived interneurons to the cerebral cortex is largely completed before birth, but subsets of inhibitory neurons, such as in the Arc pathway at the anterior lateral ventricle, may continue to migrate during infancy^10^. The diversity of GEs and GE-derived cells has been profiled in the human^11^, non-human primates^12^, and mice^13^ using single-cell transcriptomics. However, the gene regulatory and epigenomic programs that determine the regional specialization and fate of LGE, MGE, and CGE, notably between mid-gestation and infancy, are not fully defined. Furthermore, the trajectories of epigenomic and 3D genomic dynamics associated with the maturation of GE-derived neurons, especially in the basal ganglia, remain only partially characterized.

Previous studies have identified dynamic trajectories of epigenome and 3D genome conformation during neural differentiation and maturation^14–17^. These changes include the accumulation of neuronal-specific non-CG or CH cytosine methylation (mCH, H=A, T, C), which starts before birth (35-39 GW) and continues into young adulthood^15,17,18^. In the developing human frontal cortex and hippocampus, the differentiation of neural progenitor cells into neurons is associated with enrichment of short-range (SE) chromatin interactions (200kb - 2Mb), whereas differentiation into glial cells is accompanied by an increase in long-range (LE) chromatin interactions (20Mb - 100Mb)^15,16,19^. However, an increase in ultra-long-range chromatin contacts is observed during the maturation of cerebellar granule cells, the most abundant neuronal type in the cerebellum, and has been speculated to be related to the small size of granule cell nuclei^16^. The correlation between SE chromatin conformation and neuronal maturation is conserved in many mouse neuronal types, with cerebellar granular cells being a notable exception^20^. The transition to SE conformation in most types of post-mitotic neurons may impact gene regulation by increasing chromatin loops that mediate enhancer-promoter contacts and strengthening chromatin domain boundaries^15^. However, previous studies have not focused on GE or GE-derived cells, nor have they sampled primary human developing basal ganglia samples in the perinatal period. Our study thus bridges a critical gap in studying the 3D multi-omic dynamics of human brain development.

Here, we profiled the remodeling of the DNA methylome and three-dimensional genomic conformation in the GEs, developing striatum (STR), and multiple migratory destinations of CGE- and MGE-derived interneurons, including the hippocampus (HIP), and multiple cortical regions using single-nucleus methyl-3C sequencing (snm3C-seq) ^21^, spatial transcriptomics, and multimodal chromatin and RNA imaging^22^. We generated snm3C-seq profiles from over 100 primary brain specimens from 9 donors across mid-gestational, late-gestational, and infant donors. The snm3C-seq dataset is complemented by spatial transcriptomics using a highly multiplexed panel (CosMx) targeting 6,175 transcripts^23^, and a multimodal chromatin+RNA MERFISH (Multiplexed Error-Robust Fluorescence In Situ Hybridization) approach that can detect fine-scale chromatin interactions in single nuclei^22,24^. Using the dataset, we compared the multi-omic dynamics of GE-derived cellular trajectories and reconstructed regulatory programs using cellular trajectory- and branching-specific methylation states, Hidden Markov Models (HMMs)- based annotation of genomic regions^25^, and regulatory network reconstruction using DREM (Dynamic Regulatory Events Miner)^26^. Lastly, we mapped the polygenic risk for neuropsychiatric disorders to single-cell methylome profiles using met-scDRS to determine disease risk at the individual-cell level^27^.

## Results

### 3D-Epigenomic atlas of developing basal ganglia and inhibitory interneurons

We generated 155,225 new snm3C-seq profiles sampled from nine developing brains, including at mid-gestation (22-23 GW, also referred to as the second trimester or 2T), late-gestation (35, 42 GW, also referred to as 3T), 1-month post-natal (35, 42 post-natal days), and 7 months post-natal (212, 225 post-natal days) (**Figure 1A-B, Table S1**). Guided by cytological landmarks, we dissected major ventral structures, including the GEs, putamen (Pu), caudate (Ca), globus pallidus (GP), in addition to the dorsal lateral ventricle (LV). We also included major destinations of inhibitory interneuron migration, including prefrontal (DFC), frontal (FCx), temporal (TCx), cingulate (CCx) cortices, insular (ICx) and postcentral (PoCG) gyri, the hippocampus (HIP), subiculum (S), thalamus (THM), and corpus callosum (cc) (**Figure 1C, Table S1**). In addition, we integrated snm3C-seq profiles of 63,873 nuclei from our previously published developing HIP and DFC dataset generated from independent donors^15^, and 68,064 nuclei from adult human brain specimens that matched the brain regions in this study^19^. All together, we assembled an integrated dataset containing 287,147 cells spanning the complete developmental trajectory of ventral GABAergic neurons from mid-gestation progenitors to mature adult neurons.

**Figure 1.**
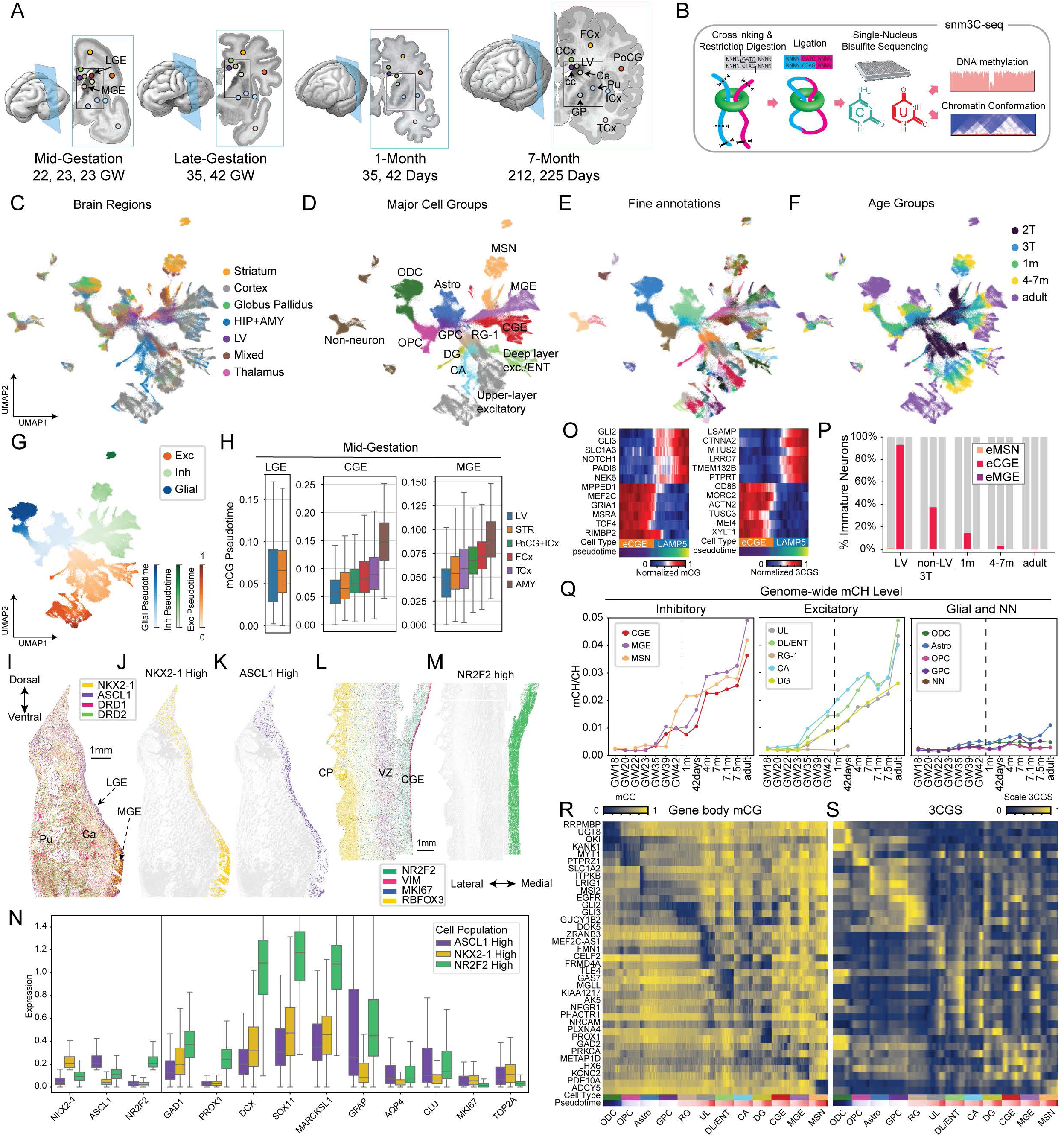
A single-cell 3D multi-omic atlas of developing human basal ganglia and inhibitory neurons. (A) The sampling of the basal ganglia and cortical regions in mid-gestational, late-gestational, 1-month postnatal, and 7-month postnatal brains. (B) Schematics of the snm3C-seq assay. (C-F) UMAP dimensionality reduction of the snm3C-seq dataset labelled with brain regions (C), major cell groups (D), fine annotations (E), and age groups (F). (G) Pseudotime scores were separately computed for excitatory, inhibitory, and glial trajectories, jointly using mCG and 3C information. (H) Distributions of single-cell mCG pseudotime scores for GE-derived neurons across sampled brain regions. (I) An anterior coronal section containing the striatal and GE regions of a 24 GW brain was labeled for the transcript abundance of NKX2-1, ASCL1, DRD1, and DRD2. (J-K) Identification of cells with high NKX2-1 (J), or high ASCL1 (K) transcript levels. (L) A posterior coronal section containing the CGE and cortical regions of a 24 GW brain was labeled for the transcript abundance of NR2F2, VIM, MKI67, and RBFOX3. (M) The selection of cells with high NR2F2 transcript level. (N) Quantification of cell identity and proliferation markers in cells with high NKX2-1, ASCL1, or NR2F2 transcript levels. (O) Normalized mCG levels (left) and 3CGS (right) of marker genes distinguishing immature CGE-derived (eCGE) and differentiated CGE-LAMP5 cells. (P) Fraction of immature neurons (eMSN, eCGE, and eMGE) in all neurons derived from each GE across developmental stages. (Q) Genome-wide accumulation of mCH during the differentiation and maturation of inhibitory (left), excitatory (mid), and glial cells (right). (R-S) Correlation of gene-body mCG (R) and 3CGS (S) at representative cell-type marker genes across single cells, organized by pseudotime scores.

The quality of the newly generated snm3C-seq dataset is similar to our previous study (**Figure S1A-C**)^15^. On average, each snm3C-seq methylome profile contains 2.4 million DNA methylome reads covering 4.24% of cytosines in the human genome (**Figure S1A-B**). Bisulfite conversion efficiency was consistently high across cell types (**Figure S1C**). After filtering out ENCODE excluded regions^28^, each snm3C-seq 3C (Chromatin Conformation Capture) profile contains an average of 214,408 chromatin contacts (136,690 intrachromosomal and 67,387 interchromosomal), with an average intra/inter chromosomal ratio of 2.0 (**Figure S1D**). Using Leiden clustering that jointly incorporates CG methylation (mCG) and 3C features^29^, we hierarchically classified the cell populations at three levels of resolution. The level 1 (L1) annotation contained four cell classes, including excitatory (Exc), inhibitory (Inh), glial, and non-neuron (NN) cells, while the level 2 (L2) annotation had 10 major groups (**Figure 1C-D, Table S2**). Among the 10 L2 groups, 9 are neural progenitor-derived. The inhibitory L1 class includes LGE-derived MSNs, as well as CGE-derived and MGE-derived interneurons. The excitatory class includes upper (UL) and deep-layer (DL) pyramidal cells, as well as hippocampal glutamatergic neurons in the cornu ammonis (CA) and dentate gyrus (DG). The glial class includes astrocytes (Astro), oligodendrocytes (ODC), and oligodendrocyte progenitors (OPC). The non-neurons (NN) include pericytes (PC), endothelial cells (EC), vascular leptomeningeal cells (VLMC), and microglia (MGC). We further clustered each major group into 225 cell types (L3 annotation) with distinct epigenomic profiles across developmental stages (**Figure 1E-F**).

### Heterogeneous developmental progressions of the ganglionic eminences

Previous single-cell analyses focused on immature inhibitory interneurons sampled in their destinations in the DFC and HIP^15^. To map the maturation trajectory of MSNs and inhibitory interneurons, we sampled the transient progenitor zones in the pre-natal GEs (**Table S1**). At mid-gestation, the anatomical proximity of inhibitory (GE) and excitatory (VZ) progenitor zones resulted in overlap during dissection. Accordingly, cells derived from both zones were combined and labeled as lateral ventricle (LV)-derived cells. We observed a major transition in the identity of LV-derived cells between mid- (2T) and late-gestation (3T) (**Figure S1E-J**), consistent with a shift from neurogenic to gliogenic activities at this developmental stage. By computing pseudotime scores separately for each of the excitatory, inhibitory, and glial trajectories (**Figure 1G**), we found that during mid-gestational brains, CGE-and MGE-derived interneurons sampled from LV had significantly lower pseudotime, corresponding to a more immature state, than inhibitory interneurons from other brain regions. This pattern was consistently observed in both mCG and 3C modalities (**Figure 1H and S1K;** Wilcoxon rank sum test, CGE-mCG, p=1.0 x 10^-105^; MGE-mCG, p=8.824 x 10^-184^). In contrast, the difference between LGE-derived MSNs sampled from LV and those from mid-gestational striatum was less significant (Wilcoxon rank sum test, mCG pseudotime p=0.008), suggesting fewer developmental differences between the cells at the LGE and immature MSNs by this stage (**Figure 1H and S1K**). The sampling of LV also extended developmental trajectories of dorsal progenitor-derived excitatory neurons. Among the mid-gestation and late-gestation samples, the LV regional dissections represent the lowest pseudotime distributions relative to cortical excitatory neurons dissected from other regions (rank sum test, mCG pseudotime p=5.2 x 10^-47^, **Figure S1L**). These results highlight LGE as a territory with fewer immature neurons relative to other neurogenic zones at 22-23 GW.

We analyzed the spatial transcriptomic profiles of the GE regions to determine neurogenic activity in 24 GW. The large, 6,175-transcript panel of the CosMx enabled the interrogation of diverse markers of cell types and states. In the anterior brain section that contains MGE and LGE regions, MGE was defined by a higher expression of NKX2-1 in the ASCL1-expressing territory (**Figure 1I-J and S1M**), whereas the LGE region was defined by less NKX2-1 expression among ASCL1+ cells (**Figure 1I-K and S1M**). On a more posterior section, CGE cells were defined by high NR2F2 expression (**Figure 1L-M and S1N**). CGE cells showed the highest expression of immature neuron markers DCX, SOX11, MARCKSL1, and pan-GABAergic neuronal marker GAD1, followed by MGE cells (**Figure 1N**). These neuronal markers were expressed at the lowest levels in LGE cells (**Figure 1N**). MGE also exhibited a higher cellular proliferation activity, as indicated by MKI67 and TOP2A expression, and a lower expression of astrocyte markers GFAP, AQP4, and CLU than in LGE (**Figure 1N and S1O-P**). Immunohistochemistry confirmed higher astrocyte marker expression of astrocyte markers S100B and GFAP in LGE territory compared with MGE (**Figure S1Q-R**). By contrast, Ki67-positive cells were most abundant in MGE, particularly at the ventricular wall. Together with the single-cell epigenomic data, these results suggest that CGE, MGE, and LGE develop at distinct rates, with the highest abundance of immature neurons in CGE, intermediate levels in MGE, and the lowest in LGE.

Lastly, we quantified immature neurons derived from each GE and found that 40% of CGE-derived cells were classified as immature neurons in late-gestational brains, compared to 0.07% and 1% for MGE- and LGE-derived cells, respectively (**Figure 1O-P**). Immature and mature neurons are robustly distinguished in clustering analysis and by mCG states and 3C Gene Score (3CGS, total chromatin interaction within a gene body) at key developmental genes (**Figure 1O-P and S1S-T**). Notably, the vast majority (93%) of CGE-derived neurons found in the LV in late gestation are immature neurons, compared to 37.4% of all CGE-derived neurons in non-LV regions, suggesting that CGE contains abundant immature neurons in late-gestation. Furthermore, CGE-derived immature neurons are the only type of immature neurons detected in post-natal brains at 1 month (14.2% of all CGE-derived cells) or at 4-7 months (2.4%) (**Figure 1P**). These results highlight the protracted developmental progression and maturation of CGE-derived neurons through infancy.

### Developmental trajectories of lineage-specific DNA methylation

Neurons and glial cells in the human brain have high global levels of mCG throughout the lifespan, whereas mCH accumulates specifically in neurons and constitutes a unique feature of the neuronal DNA methylome (**Figure 1Q and S1U**)^17^. mCH begins to accumulate in neurons during late gestation and continues to increase as neurons mature during childhood^15,17,18^. The pattern of mCH accumulation is specific to each major lineage of neurons (**Figure 1Q**). Inhibitory CGE and MGE lineages have similar patterns of mCH accumulation starting late in the third trimester (between 35 and 39 GW). LGE-derived MSNs follow a distinct trajectory, rapidly accumulating mCH during perinatal development between 39 GW and 1-month after birth. The level of mCH in mature LGE-derived cells is between that of CGE and MGE-derived cells (**Figure 1Q**). Consistent with previous observations^15^, excitatory CA neurons of the hippocampus have the earliest onset of mCH, beginning in GW 35. Excitatory DG neurons have the lowest levels of mCH relative to other excitatory lineages (**Figure 1Q**). Glial cells accumulate much less mCH than neuronal types.

The functional significance of DNA methylation is suggested by the strong correlation of high mCG and mCH with low mRNA expression^17,19,30^. Consistent with this, we observed a significant negative correlation between gene body mCG and 3CGS (**Figure S1V**). Gene body mCG and 3CGS are strongly anti-correlated at canonical cell type marker genes (**Figure 1R-S**).

### Distinct development trajectory of LGE-derived MSNs

We determined the developmental trajectory of LGE, CGE, and MGE-derived cell populations across stages, using the unbiased clustering of genome-wide mCG patterns and gene body mCG at key marker genes (**Figure 2A and S2A-B**). In contrast to CGE-and MGE-derived cells, whose subtypes (e.g., MGE-PVALB) are not distinguishable until late gestation (**Figure S2A-B**), all LGE-derived MSN subtypes were identifiable in mid-gestation. In addition, we identified an immature MSN (eMSN) population that is only found in mid-gestational samples, and which may give rise to more differentiated MSN subpopulations (**Figure 2A and S1S-T**). MSN subtypes were primarily distinguished by mCG signatures (**Figure 2B-D**) and, to a much lesser extent, by 3C contact features (**Figure S2C-D**). The UMAP embedding of developing MSNs revealed distinct cell populations, with few intermediate cells between ages and MSN subtypes, in contrast to the continuous pattern observed in all other types of neurons (i.e., cortical and hippocampal excitatory neurons and inhibitory interneurons) characterized in this study (**Figure 2B-D and 1D**). Each of the main MSN classes (DRD1 and DRD2) was separated into matrix (EPHA4+) and striosome (BACH2+) subpopulations (**Figure 2C**), and these were further organized into putamen- and caudate-localized populations (**Figure 2D**). In addition, eccentric MSNs exhibit greater differences from matrix or striosome MSNs across ages and dissected brain regions (**Figure 2C,E, and S2E-F**).

**Figure 2.**
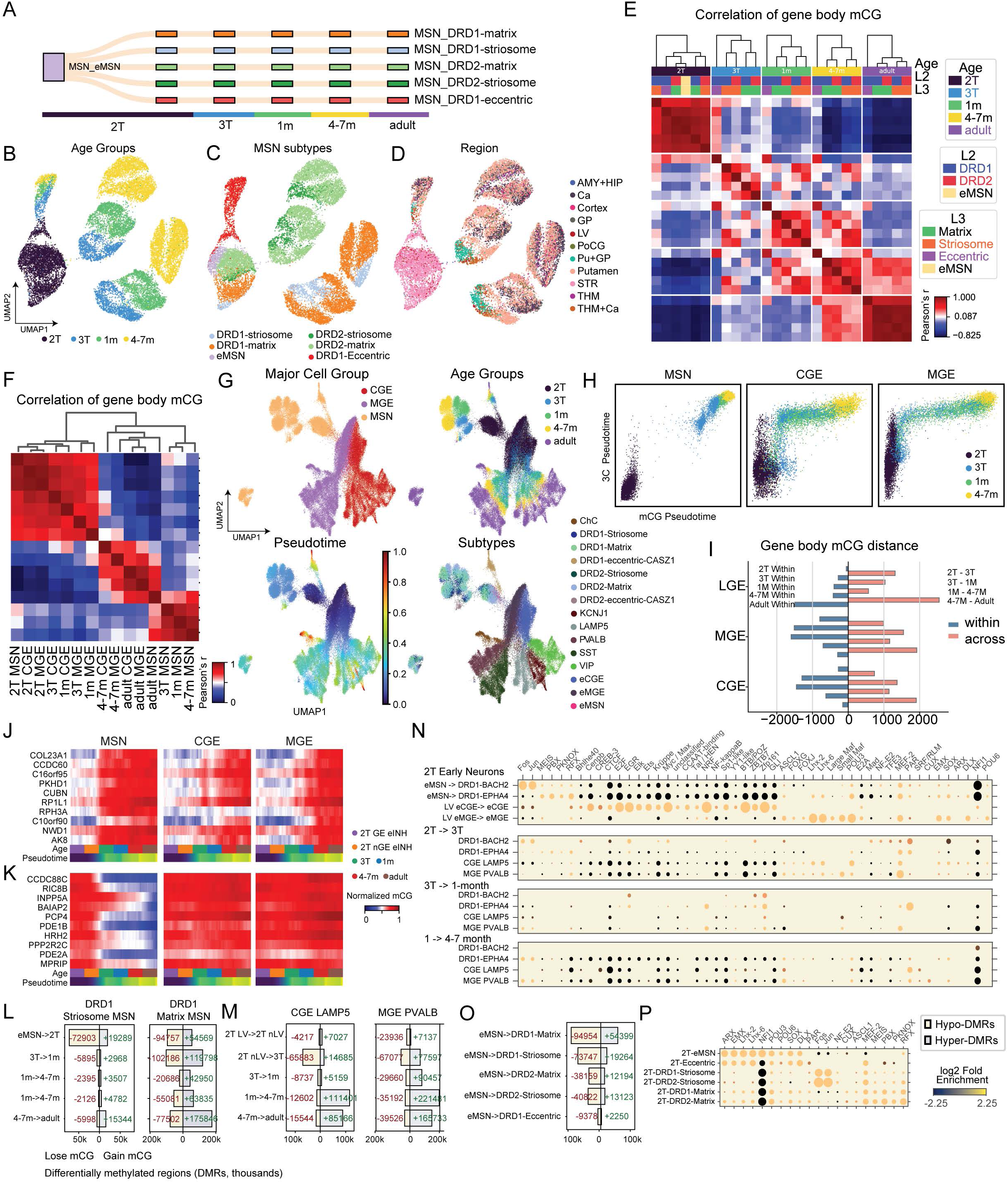
Distinct developmental trajectory and regulatory signatures in LGE-derived MSNs. (A) The specification of MSN subtypes in mid-gestation. (B-D) UMAP dimensionality reduction of MSN mCG profiles labelled with age groups (B), subtypes (C), and brain regions (D). (E) Correlation matrix of MSN subtypes across age groups computed using scaled gene body mCG. (F) Correlation matrix of GE-derived inhibitory neurons across age groups computed using scaled gene body mCG. (G) UMAP dimensionality reduction of all GE-derived inhibitory neurons labelled with GE regions (upper left), age groups (upper right), pseudotime scores (lower left), and subtypes (lower right). (H) Comparison of pseudotime scores computed using mCG and 3C modalities across the differentiation of LGE-derived MSNs, and CGE- and MGE-derived interneurons. (I) Quantification of intra-cell-type and inter-cell-type gene body mCG distances in LGE, CGE, and MGE-derived cell populations. (J-K) Scaled gene body mCG of selected marker genes during the differentiation of LGE, CGE, and MGE-derived cell populations. The marker genes were selected for demethylation (J) and gain of methylation (K) during MSN differentiation. (L) Trajectory-DMRs identified across the differentiation of DRD1-Striosome, DRD1-Matrix. (M) Trajectory-DMRs identified across the differentiation of CGE-LAMP5, and MGE-PVALB. (N) TF binding motif enrichments in trajectory hypo-DMRs. (O-P) Branch-DMRs (O) and the associated TF binding motif enrichment (P) identified during the specification of MSN subtypes in mid-gestation.

We observed a hierarchical organization of the developing and mature MSN subpopulations using gene body mCG (**Figure 2E**). MSN subtypes clustered primarily by developmental stage, followed by local tissue context (matrix vs. striosome vs. eccentric), then dopamine receptor class (DRD1 vs. DRD2), and finally by striatal region (caudate vs. putamen) (**Figure 2E, S2E-F**). The DNA methylome of immature MSNs in mid-gestation closely reassembled developing MGE and CGE neurons, but took on a unique configuration by late gestation, which persisted following birth (**Figure 2F**). These data show that MSNs rapidly differentiate between mid- to late-gestation and between infancy and adulthood.

We quantified the epigenomic similarity between cells derived from the GE regions, using gene body mCG levels, and found greater similarities between CGE- and MGE-derived neurons, while MSNs were more distinct (**Figure S2G-J**). UMAP embedding of GE-derived cells based on mCG and 3C features revealed a more discrete distribution of MSNs across age groups compared to CGE- and MGE-derived inhibitory interneurons (**Figure 2G**). Notably, pseudotime analysis of mCG and 3C modalities revealed a two-step sequence in the development of inhibitory interneurons derived from the MGE and CGE (**Figure 2H**). In these cells, the chromatin interaction landscape (3C pseudotime) undergoes a major wave of maturation in mid- and late-gestation, followed by maturation of the mCG landscape in late-gestation (3T) and post-natally (**Figure 2H**). By contrast, our data from developing MSN showed no sequential relationship between 3C and mCG maturation. Instead, both modalities showed a major transition between 2T and 3T samples (**Figure 2H and S2K-M**), consistent with the correlation analysis **(Fig. 2F)**. This suggests that DNA methylation and chromatin conformation develop synchronously or that any sequential dynamics occurs over a brief time window during late gestation (**Figure 2H and S2K-M**). Consistent with the appearance of homogeneity among MSNs, quantification of cell-to-cell distance as a total mCG change summed across all genes showed that MSNs consistently exhibit smaller within-age-group distances among individual MSN cells and greater between-age-group distances (**Figure 2I**), consistent with more synchronous maturation compared to inhibitory interneurons.

The genome-wide dynamics were corroborated by mCG dynamics at marker genes (**Figure 2J-K and S2N-O**). Genes that gain mCG during MSN development, likely indicating transcription repression in mature neurons, show sharp changes between mid- and late-gestation. By contrast, in CGE- and MGE-derived neurons, these genes remodel gradually between late-gestation and infancy (**Figure 2J**). Genes that lose mCG in MSNs are more cell-type specific, with genes selected for strong demethylation during MSN, indicating transcriptional activation, are more lineage specific: these genes largely retain high levels of mCG in CGE- and MEG-derived neurons (**Figure 2K**). The mCG dynamics at these genes support the presence of DNA methylation signatures distinguishing MSNs from interneurons, and the distinct temporal patterns of epigenomic remodeling in MSNs.

### Genomic regulation of MSN subtypes

To compare regulatory signatures between GE-derived neurons, we identified differentially methylated regions (DMRs) between succeeding developmental stages (trajectory-DMRs)^15^. We previously found that DMRs that lose methylation (hypo-DMRs) during mid-gestation are enriched for binding motifs of lineage-specific transcription factors (TFs). In contrast, DMRs that lose methylation in late gestation or early infancy are enriched for motifs of activity-dependent TFs, including AP-1 (FOS, JUN) and EGR. Here, we found a striking difference in the regulatory motifs associated with developmental DMRs in DRD1+ MSNs from the matrix (DRD1-EPHA4) and striosome (DRD1-BACH2) compartments. The majority of DMRs in DRD1-Striosome MSNs lost methylation during mid-gestation (eMSN -> 2T, **Figure 2L**), and these DMRs were strongly enriched for the binding motifs of neuronal activity-dependent AP-1 TFs **(Figure 2N)**. By contrast, DRD1-Matrix DMRs that lose methylation in mid-gestation were depleted for the AP-1 motif **(Figure 2N)**. At the same time, the binding motifs of MEIS, PBX, and PKNOX were more strongly enriched in the mid-gestational hypo-DMRs found in DRD1-Matrix than in DRD1-Striosome. Hypo-DMRs in DRD1-Matrix, as well as eccentric MSNs (**Figure S2Q-R**) were enriched in AP-1 motifs during infancy, much later than the appearance of the neuronal activity signature in DRD1-Striosome. This finding suggests that striosomal MSNs are associated with greater expression of AP-1 immediate-early genes during mid-gestation, compared to matrix MSNs or interneurons.

We found differences in regulatory signatures between hypo-DMRs in each of the three GEs. Hypo-DMRs in early immature MGE-derived neurons (eMGE) were enriched in binding site motifs of canonical MGE-defining TFs such as LHX6 and MAF (**Figure 2M-N**). By contrast, LGE-derived MSN hypo-DMRs were enriched in motifs for MEIS, PBX, and PKNOX. On the other hand, in hyper-DMRs across multiple GEs, we observed enrichment of binding motifs for early patterning TFs that mark inhibitory progenitor zones, such as SOX, ARX, and DLX, suggesting a shared signature of progenitor program termination (**Figure S2P**). The increase of mCG at hyper-DMRs occurred earliest in LGE-derived MSN neurons in mid-gestation (**Figure S2P**). This was consistent with our findings that the neurogenic activity in LGE precedes MGE and CGE, and with the onset of gliogenesis in LGE during mid-gestation (**Figure 1N and S1Q-R**).

Lastly, we identified branch-DMRs: DMRs that distinguish differentiated cell populations from the shared progenitor cells (**Figure 2O-P**). This analysis recapitulated the strong enrichment of the AP-1 motif in striosomal DRD1+ and DRD2+ neurons (**Figure 2P**). Notably, we found that eccentric MSNs remain similar to immature MSNs, with a relatively small number of DMRs (**Figure 2O**). The TF binding profile of eccentric MSNs was similar to that of eMSNs, except for a notable depletion of Nuclear Factor 1 binding motif, consistent with the transition to post-mitotic neurons (**Figure 2P**).

### 3D Genome Reorganization in MSN Subtypes

In addition to dynamic changes in DNA methylation, neural differentiation and maturation involve reorganization of the 3D configuration of DNA chromatin within the nucleus at multiple genomic scales. At the largest scale, we analyzed megabase-scale domains of active (A) or inactive (B) chromatin compartments in developing MSNs. We found substantial developmental remodeling of A/B compartments across the MSN lineage. For example, a region on chromosome 4 (chr4: ∼54 Mb) encompassing the MSN developmental regulator GSX2^31,32^ transitions from an active A compartment configuration in mid-gestation to a repressive B compartment by 4–7 months **(Figure 3A-B)**. Across developmental stages, compartment profiles clustered by MSN subtype, with striosome and eccentric MSNs clustering together independently of DRD1/DRD2 identity **(Figure 3C)**. Variance partitioning confirmed that developmental stage explains the largest fraction of variance in compartment scores, followed by matrix–striosome identity, whereas DRD1/DRD2 identity contributes relatively little **(Figure S3A)**. Regional differences between caudate and putamen MSNs were modest **(Figure S3B)**.

**Figure 3.**
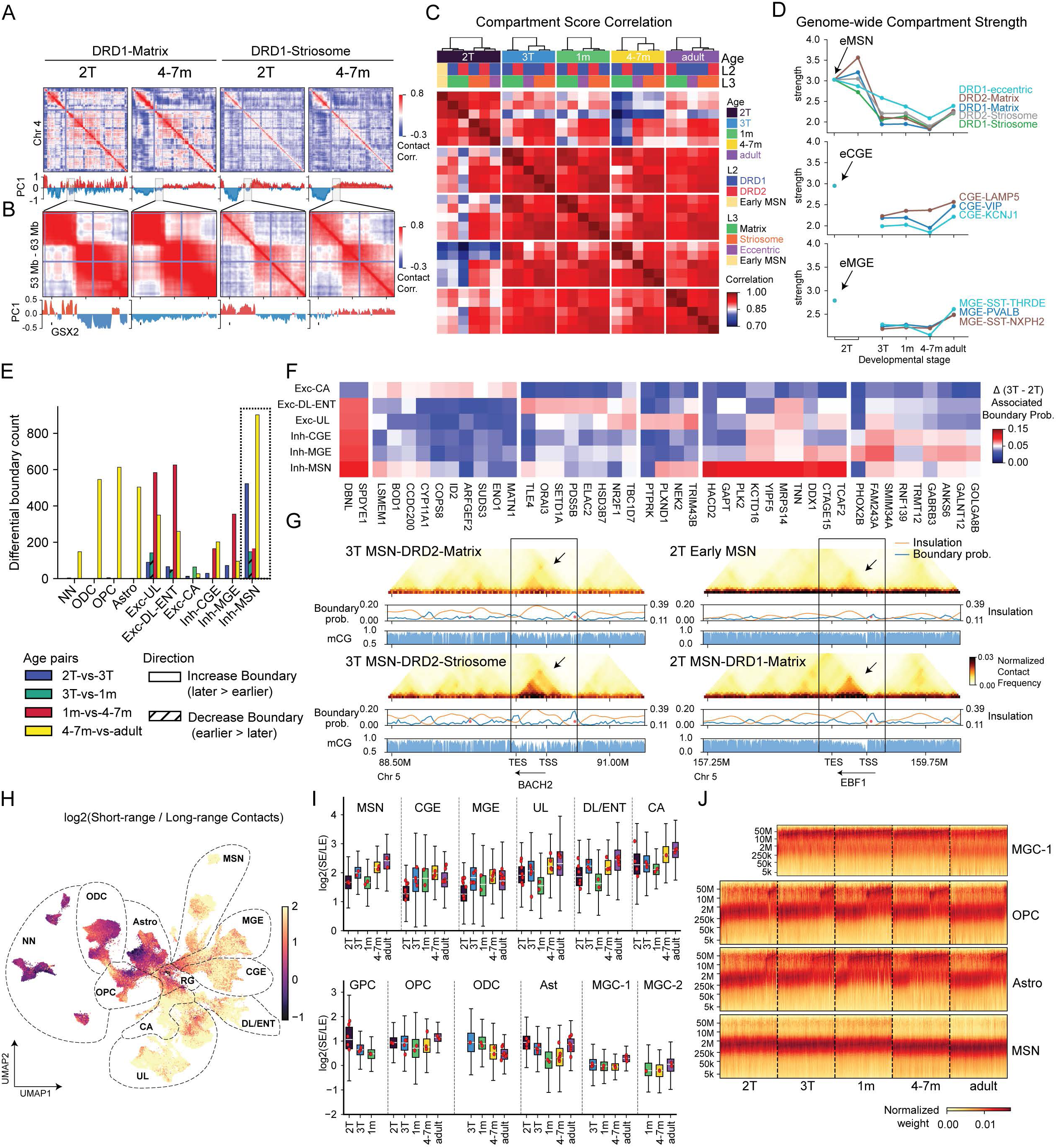
3D Genome reorganization in LGE-derived MSN. (A) Top rows show correlation matrices of chromosome 4 derived from distance-normalized contact maps for DRD1-Matrix and striosome MSNs in mid-gestation and at 4–7 months. Bottom rows show the first principal component (PC1) of the corresponding correlation matrices, where positive (red) and negative (blue) values indicate A and B compartments, respectively. (B) Same as (A), shown for a zoomed genomic region on chromosome 4 (52–67 Mb). (C) Genome-wide Pearson correlations of A/B compartment scores across MSN subtypes. (D) Compartment strength across the developmental trajectories of MSN, CGE and MGE-derived neurons. (E) Number of differential domain boundaries identified between adjacent age groups. Solid bars indicate boundaries gaining strength, whereas cross-hatched bars indicate boundaries losing strength. (F) Top-ranked genes proximal to age-related differential TAD boundaries between the second and third trimesters, selected separately for each major neuronal trajectory. Heatmaps show changes in boundary probability (3T − 2T). (G) Heatmaps on the left show normalized chromatin contact maps for DRD1-Matrix and DRD1-Striosome MSNs in the third trimester, centered on a cell-type–specific domain boundary at the BACH2 locus. Orange and blue tracks indicate TAD insulation scores and boundary probabilities, respectively, together with mCG methylation at bottom. On the right, heatmaps for early MSNs and DRD1-Matrix MSNs in the second and third trimesters highlight an age-associated domain boundary at the EBF1 locus. (H-I) Log₂ short-/long-range chromatin contact ratio shown on UMAP (H) and as boxplots (I). (J) Heatmaps show contact frequency by genomic distance in individual cells across developmental stages. Cells were subsampled to equal numbers per stage, and contact counts were normalized within each cell by total contacts across distances, enabling comparison across neuronal populations (e.g., MSNs) and glial cell types, including astrocytes, OPCs, and MGC-1 microglia.

In addition to compartment switching, we observed quantitative changes in the strength of the compartment signal, reflecting the degree of segregation between A and B compartments. Compartment strength was higher in 2T MSNs, followed by a marked reduction between 2T and 3T **(Figure 3D)**. We observed a similar trend of reduced compartment strength in developing interneuron and excitatory neuron lineages. By contrast, glial and microglial populations had a progressive increase in compartment strength during development **(Figure 3D, S3C).** Consistent with these trends, dcHiC^33^ identified more events of compartment switching (A-to-B, or B-to-A) between the second and third trimester than in any other interval in MSNs **(Figure S3D)**.

To investigate chromatin organization at a finer genomic scale relevant for individual genes, we examined topologically associated domains (TADs) using insulation scores at 25 kb resolution to quantify domain boundaries. While genome-wide insulation profiles were largely conserved across MSN subtypes and developmental stages (**Figure S3E)**, the number of chromatin domains increased progressively during development **(Figure S3F)**. To characterize these changes, we identified age-related differential boundaries (age-DBs) that capture stage-specific remodeling of chromatin domain organization (**Methods**). In MSNs, age-DBs emerged across multiple developmental windows, with many appearing between the 2T and 3T and additional changes occurring postnatally (4–7 months relative to adulthood). In contrast, interneuron (CGE- and MGE-derived) and excitatory neuron lineages showed predominantly postnatal boundary remodeling, with the largest shifts observed between 1 month and 4–7 months of age **(Figure 3E)**. The patterns mirror the global DNA methylome correlation analysis (**Figure 2F**), consistent with early prenatal differentiation of MSNs followed by more gradual maturation.

To investigate the potential functional relevance of dynamic changes in local chromatin organization, we focused on the second-to-third trimester interval and examined genes located near age-related differential TAD boundaries. In MSNs, HACD2, GAPT, and PLK2, together with numerous chromatin-associated and cellular infrastructure genes, are located proximal to MSN-specific chromatin boundaries, implicating local domain remodeling during MSN development **(Figure 3F)**. Consistent with this, we observed cell subtype-specific domain dynamics during early MSN maturation. The striosome-enriched gene BACH2 resided within a stronger chromatin domain in striosome than in matrix MSNs **(Figure 3G, left).** Moreover, the key regulatory genes of matrix MSNs, such as EBF1, progressively associated with strengthened domains during maturation **(Figure 3G, right)**.

Finally, at the fine-scale genomic scale, we examined chromatin loops and tested for differential looping across major developmental lineages **(Methods)**. Across all lineages, developmental loop gains substantially outnumbered loop losses, indicating progressive establishment of regulatory interactions during maturation **(Figure S3G-H)**. Annotating differential loops to nearby gene promoters revealed distinct functional signatures across lineages: chromatin loops in neurons were enriched for genes involved in neuronal development and synaptic function, whereas loops in glial lineages were associated with metabolic and immune-related processes **(Figure S3I)**, as determined by Gene Ontology enrichment analysis^34^. Together, these findings suggest that developmental chromatin looping reflects lineage-specific functional programs rather than a uniform regulatory trajectory.

### Transient short-range chromatin conformation is associated with OPC to ODC differentiation

Neurons, including MSNs, show iincreased short-range (SE, 200 kb–2 Mb) chromatin contact frequency relative to long-range (LE, 20–100 Mb) interactions, providing a potential structural basis for strengthened local chromatin domains^15,19,35^. To quantify these differences, we computed the log₂ ratio between short-range (200 kb–2 Mb) and long-range (20–100 Mb) contact frequencies (**Figure 3H**). Across both excitatory and inhibitory neurons, increasing domain strength is accompanied by a progressive enrichment of short-range contacts. Notably, most neuronal cell types pass through a transient state around 1 month after birth, characterized by lower short-range contact enrichment than in the third trimester of gestation and in older infants at 4–7 months **(Figure 3H-I)**.

In contrast, glial populations, including astrocytes, OPCs, and ODCs, had a striking bimodal distribution of SE and LE states across all developmental stages (**Figure 3J**). This pattern was not observed in neurons or microglia (**Figure 3J**). Notably, similar SE/LE patterns were observed across all dissected brain regions **(Figure S4A)**, indicating that this heterogeneity is unlikely to be driven solely by regional differences in cell composition. Although astrocytes also exhibited SE/LE heterogeneity, the patterns of heterogeneity are not clearly associated with age, brain regions, or astrocyte subtypes identified by methylation signatures **(Figure S4B)**.

To investigate the dynamics of SE/LE genome configurations, we focused on the OPC-to-ODC differentiation. Embedding glia progenitor cells (GPCs), OPC, and ODC in pre- and post-natal brains based on 3C features revealed a population of cells that formed a continuum between OPC and ODC states, which we termed transitional OPC (tOPC) and transitional ODC (tODC) (**Figure 4A**). These cells arose mainly from 1-month postnatal brain samples (**Figure 4B and S4C**). tOPCs were enriched at 1 month in both cortex and striatum, whereas enrichment of tODCs at this stage was observed primarily in the striatum (BH-FDR < 0.05, Fisher’s exact test; **Figure S4D**). Although we anticipated SE enrichments would be found in 2T GPC cell population due to their early stage of development, we found surprising SE enrichments in tOPC and tODC populations in infant brains, as well as aOPC and srODC cells in the adult brain (**Figure 4C-D, Figure S4E**). The tOPC-tODC transition also showed the most rapid change in 3C pseudotime (Figure 4E), whereas mCG pseudotime was less dynamic (**Figure 4F**). These features observed during the OPC-ODC transition indicate major 3D genomic rearrangements during OPC differentiation to ODC.

**Figure 4.**
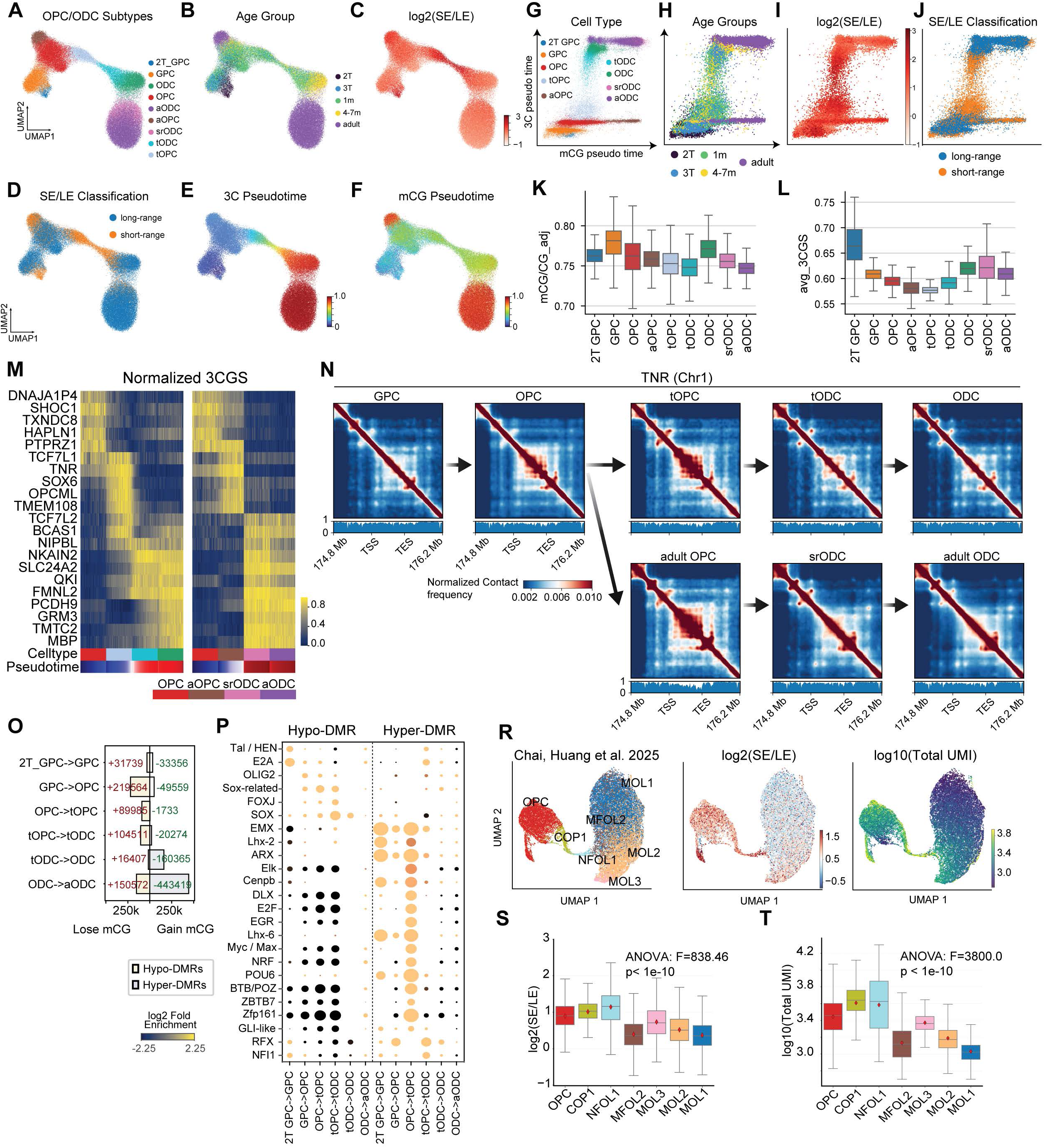
Transient global chromatin conformation remodeling during the differentiation of OPC to ODC. (A-F) UMAP dimensionality reduction of the 3C modality for OPC and ODC cells labeled with subtypes (A), age groups (B), log₂(short/long) range interactions (C), SE/LE classification (D), 3C pseudotime (E), and mCG pseudotime (F). (G-J) Scatter plots comparison of mCG and 3C pseudotime scores labeled with subtypes (G), age groups (H), log₂(short/long) range interactions (I), and SE/LE classification (J). (K) Genome-wide methylation changes across subtypes. (L) Global reduction of 3CGS during the transition between tOPC and tODC. (M) Genes showing 3CGS dynamics during transition from tOPC to tODC. (N) Local chromatin confirmation dynamics at TNR during the differentiation of ODC in infant and adult brains. (O) Number of trajectory-DMRs and TF binding motifs (U) identified across the differentiation of ODC in developing brains. (R) UMAP embedding of the transcriptomic profiles of mouse OPC and ODC profiled using ChAIR^20^, labeled with cellular subtypes^20^ (left), log₂(short-range / long-range) chromatin interactions (mid), and log₁₀ total UMI counts as a proxy for genome-wide transcriptional activity (right). (S-T) Quantification of log₂(short-range / long-range) chromatin interactions (S) and log10(total UMI counts) for each cell profiled in the ChAIR dataset (T).

Joint analysis of mCG and 3C pseudotime across the OPC-ODC differentiation trajectory revealed complex, multi-modal dynamics (**Figure 4G-J**). Early differentiation from glial progenitor cells (GPCs) to OPCs was followed by a bifurcation of OPC to ODC differentiation in late gestation and infancy. At this stage, a major branch of cells progressed toward mature ODCs, with major changes in chromatin conformation. A second, smaller subset of OPCs exhibited maturation trajectories towards adult OPCs (aOPCs), with increased DNA methylation pseudotime but little change in 3C pseudotime (**Figure 4G-H**). Notably, the majority of cells with intermediate 3C pseudotime, between the OPC and ODC states, were found during infancy. Although we expect that a fraction of adult OPCs would continue to differentiate into ODC, we detected few intermediate cells despite extensive sampling of adult cells (27,160 OPCs or ODCs) (**Figure 4H**). SE chromatin organization was enriched in transitional tOPC and tODC populations **(Figure 4J)**, indicating the close association between chromatin state and progression along OPC-ODC lineage.

Chromatin remodeling during the transition from OPC to ODC is correlated with a moderate reduction of mCG (**Figure 4K**) and 3CGS **(Figure 4L)**, consistent with the observation that short-range contacts are less frequently associated with intragenic regions **(Figure S4F-G)**. Notably, this remodeling was particularly evident at the TNR locus **(Figure 4M-N)**, a key regulator of oligodendrocyte maturation and axon–glial interaction ^36^. TNR exhibited a transient increase in 3CGS and an elevated number of chromatin loops in tOPC, consistent with dynamic regulatory reorganization during the OPC-to-ODC transition. Similarly, other chromatin domain dynamics were also observed at other lineage-associated loci, including OPCML **(Figure S4H)** and SOX6^37^ **(Figure S4I)**, across both infant and adult ODC differentiation trajectories. We identified DMRs and TF binding motifs along the OPC to ODC differentiation trajectories in infant (**Figure 4O-P**) and adult (**Figure S4J-K**) brains. The results recapitulate known transcriptional regulations, including the enrichment of OLIG2 motif at the specification of OPC, and the enrichment of SOX motif during the transition between tOPC and tODC in the infant brain (**Figure 4P**), and the transition between aOPC and srODC in the adult brain (**Figure S4K**).

We previously found that the SE chromatin conformation is associated with an increase in chromatin loops and domain boundaries^15^, implicating more active genome-wide transcription activities. To explore the functional implications of transient enrichments in short-range chromatin interactions, we analyzed a recently published tri-omic ChAIR dataset^20^ from developing adult mouse brains, which jointly profiles chromatin accessibility, transcription, and chromatin conformation. Consistent with our findings in the human brain, we observed that cells at the transitional stages, including committed oligodendrocyte precursor (COP1) and newly formed oligodendrocyte (NFOL1), are strongly enriched for short-range chromatin contacts, distinguishing them from both upstream OPCs and downstream ODCs (**Figure 4R**). Leveraging matched transcriptomic measurements, we further found that these transitional cells exhibit elevated transcriptional activity, as reflected by a higher number of total unique mRNA molecules detected (**Figure 4S-T**). This result supports the association between SE and greater genome-wide transcriptional activity, which may facilitate the transition of OPCs to ODCs.

### Spatial mapping of developing basal ganglia cell types

We integrated snm3C-seq with CosMx spatial transcriptomic profiles to determine the relationships between nuclear area, transcriptional activity, and genome-wide chromatin conformation in the developing basal ganglia (**Figure 5A-D and S5A-B**). In both 24 GW and 35 GW basal ganglia samples, cell populations annotated via integration with snm3C-seq references show expected expression patterns of cell-type-specific markers (**Figure S5C-D**). Mapping the spatial location of annotated cell types revealed their characteristic locations (**Figure 5E-H and S5G-H**), such as DRD1-Matrix and DRD1-Striosome localizing in complementary compartments (**Figure 5E-H**), the location of immature neurons eMSN and eMGE in the GE in the 24 GW section (**Figure S5G**), or the enrichment of astrocytes in the ventricle zone in the 35 GW section (**Figure S5H**). The spatial mapping of cell populations corroborates the heterogeneous maturation progress of GEs. Immature eMGE cells substantially outnumber eMSN cells in the progenitor zone in the 24 GW section, while astrocytes are more abundant in LGE than in MGE regions (**Figure S5G**). The nuclear area of all annotated cell types increases between 24 and 35 GW (**Figure 5I and S5I-J**). The nuclei of striosome MSNs are significantly larger than matrix MSNs in 24 GW, whereas the differences diminish in 35 GW (**Figure 5I**), consistent with the earlier maturation of striosome MSNs. In line with the finding in the adult mouse brain^38^, we found strong correlations between nuclear area and total transcript counts in the developing human basal ganglia (24 GW: Pearson’s r = 0.796, p = 1.1 x 10^-3^; 35GW: r= 0.885 for 35 GW, p = 2.6 x 10^-5^) (**Figure 5J and S5K**).

**Figure 5.**
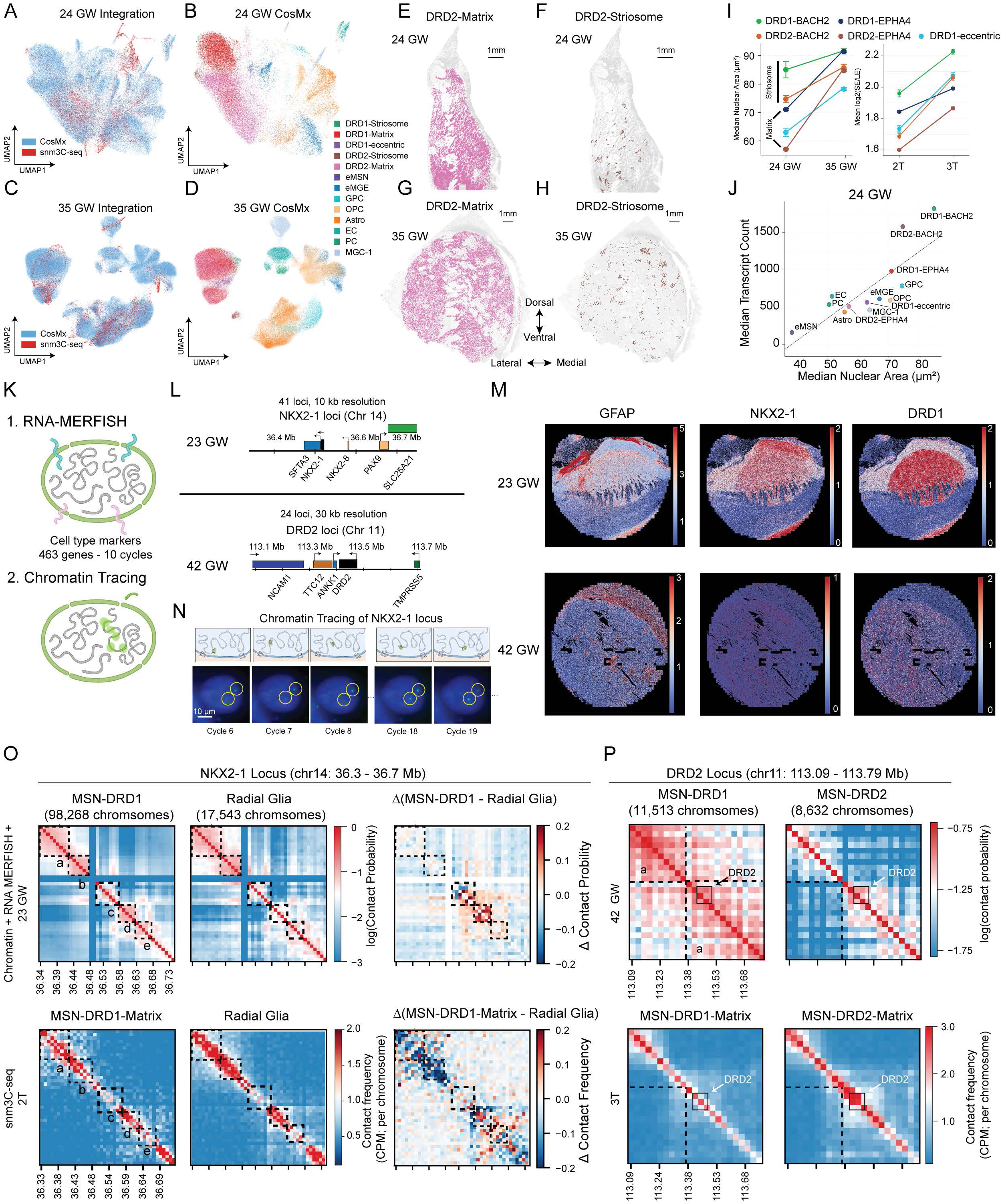
Spatial mapping of developing basal ganglia cell types. (A-D) Integration of snm3C-seq with CosMx spatial transcriptomic profiles generated from 24 GW (A-B), and 35 GW (C-D) sections containing the developing basal ganglia. (E-H) Spatial localization of matrix (E) and striosome (F) DRD2-expressing neurons in the 24 GW section, and matrix (G) and striosome (H) DRD2-expressing neurons in the 35 GW section. (I) Estimated nuclear area increased from 24 to 35 GW across annotated cell types (left), and were correlated with the ratio between short and long range interactions (right). (J) Correlation between nuclear area and total transcripts count in the 24 GW section. (K) Schematics of chromatin tracing followed by RNA MERFISH+. (L) Genomic design and probe coverage for chromatin tracing at the NXK2-1 and DRD2 loci. (M) Spatial maps of maker-gene expression in the 23 GW and 42 GW samples. (N) Representative chromatin tracing images. (O) Comparison of chromatin tracing and snm3C-seq profiles at the NKX2-1 locus in the 24 GW sample for MSN-DRD1 and radial glia. (P) Comparison of chromatin tracing and snm3C-seq profiles at the DRD2 locus in the 42 GW sample for MSN-DRD1 and MSN-DRD2.

To directly interrogate the 3D organization of key developmental loci in specific cell types in situ, we performed imaging-based chromatin tracing ^39^ alongside a 463-gene RNA MERFISH+ panel to measure spatial distances between genomic segments and assign cell types **(Figure 5K)**. Our DNA probes targeted two loci: (i) the NKX2-1 locus in mid-gestational tissue and (ii) the DRD2 locus in late-gestational tissue. We designed sequential DNA FISH probe sets spanning each locus at 10-kb/30-kb resolution **(Figure 5L)**. In parallel, MERFISH profiling of the transcript panel enabled assignment of major cell classes in both 23 GW and 42 GW samples **(Figure 5M and S5L)**, and chromatin tracing enabled reconstruction of 3D conformations at the single-cell level **(Figure 5N)**. At the NKX2-1 locus in mid-gestation, chromatin tracing revealed pronounced domain structures and clear differences between MSN-D1 cells and radial glia **(Fig. 5O top panels, S5M,N)**. The local domain structures were less resolved in 3C profiles (**Figure 5O lower panels**). In contrast, we found that 10kb-resolution chromatin tracing revealed fine-scale interaction features, such as foci a-e in **Figure 5O and S5N**, that were not apparent in the corresponding 3C maps. At the DRD2 locus in late gestation (42 GW), chromatin tracing further showed notable domain separation in DRD2-expressing MSN subtypes that was not observed in DRD1-expressing MSN subtypes (**Figure 5P and S5O**). However, such a cell-type-specific pattern was not detected in 3C contact maps, which indicate comparable TAD separation in both DRD1-and DRD2-expressing MSN subtypes **(Fig 5P, S5O)**. As an internal control, VLMC exhibited consistently weak domain structure in both chromatin tracing and 3C contact maps (**Fig S5O)**. This discrepancy suggests that the imaging-based chromatin tracing and proximity-ligation-based 3C may differ in their ability to detect certain locus and cell-type-specific chromatin interactions, highlighting the complementary utility of these two types of approaches.

### Cortical regionally specific epigenomic patterns

The incredible diversity of excitatory neurons that form the cortical layers arises from RG cells in the VZ-SVZ (**Figure 6A**)^40^, differentiating to intermediate progenitor cells and later differentiated excitatory neurons (**Figure 6A and S1L**). Our multi-regional dissection of progenitor zones shows that 82% of cell samples from VZ are RG cells (**Figure 6B**). By including 1-month-old cases, we found that the vast majority of excitatory cell types identified in the adult brain can already be distinguished by mCG signatures at postnatal 1-month (**Figure 6A**). Gene body mCG remodeling is enriched in mid- to late gestation among genes showing the strongest changes in mCG (**Figure 6C-D**), supporting the establishment of cell-type-specific methylation patterns before birth.

**Figure 6.**
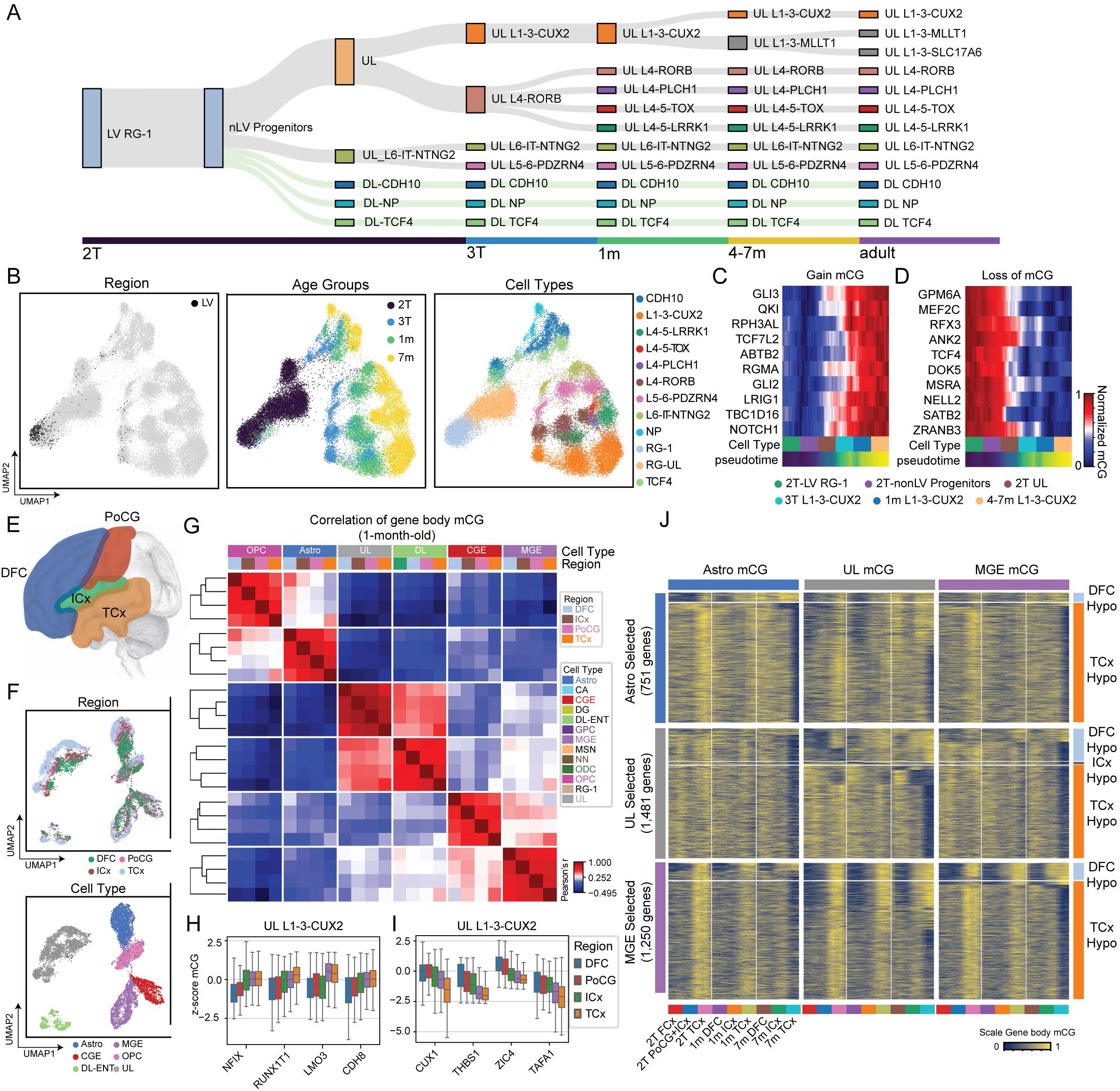
Excitatory cell type regionally specific epigenomics. (A) The differentiation of cortical excitatory neuronal cell types from VZ-SVZ in mid-gestation to layered cortical subtypes in adults. (B) UMAP embedding of cortical excitatory trajectories including cells derived from mid-gestation to 7 months donors, labeled with all cells derived from dissections targeting LV ((left), age groups (middle), and cell types (right). (C-D) Genes selected for the strongest positive (C) or negative (D) correlations between gene body mCG to L1-3-CUX2 trajectory pseudotimes. (E) Schematic of cortical areas. (F) mCG UMAP embedding of cortical cells derived from donor 6419 (1-month-old) colored by cortical regions (top) and cell types (bottom). (G) Pearson correlation of gene body mCG in major cell groups across DFC, ICx, PoCG, and TCx. (H-I) Z-scaled mCG for select genes showing variable gene body mCG across cortical regions in multiple major cell groups. (J) Genes exhibiting cortical-regional-specific methylation signatures were identified in astrocytes, upper-layer neurons, and MGE-derived neurons.

Transcriptomic profiling has revealed that across the many functionally distinct regions of the cortex, the same excitatory cell types are present but exhibit region-specific characteristics^41^. To remove the donor effect on the relatively moderate cortical regional differences, we analyzed the methylome signature of cortical arealization within a single 1-month-old donor, 6419 (**Figure 6E-G**), and replicated in a 23 GW donor **(Figure S6A-B),** and a 7-month-old donor **(Figure S6C-D)**. Cortical regional differences in the methylome are reproducibly detected in multiple cell types (**Figure 6E-F**) and are consistent across ages (**Figure S6A-D**). The DFC-TCx difference is the greatest among all pairwise comparisons of cortical regions in all major cell groups with significant cortical representation **(Figure 6G and S6B,D)**. In upper layer excitatory neurons (UL L1-3-CUX2), the top genes whose gene mCG patterns show a significant correlation to cortical regions are transcriptional regulators, including NFIX, RUNX1T1, LMO3, CUX1, ZIC4, and genes with cell adhesion and extracellular matrix functions, including CDH8, THBS1, and TAFA1, and presumably modulate cellular migration or axon guidance (**Figure 6H-I**). We identified 751, 1,481, and 1,250 genes associated with cortical-regional methylation signatures in astrocytes, upper-layer excitatory neurons, and MGE-derived inhibitory interneurons, respectively (**Figure 6J and Table S3**). Strikingly, genes showing hypomethylation in TCx account for the vast majority (94.8% (712/751) in Astro, 72.2% (1,069/1,481) in UL, and 89.2% (1,115/1,250) in MGE) of the signature genes (**Figure 6J, S6E-G, and Table S3**). In addition, TCx-specific hypomethylation strengthens across development and becomes more pronounced in the 4-7-month-old brain (**Figure 6J**). Gene ontology analysis of the cortical-regional genes identified transcription regulation being the most strongly enriched terms, followed by neurogenesis-related terms.

### ChromHMM-based genomic annotation using combinatorial methylation signatures

To further identify and characterize the global patterns of methylation across cell types and age groups, we utilized ChromHMM^25^. While ChromHMM is most commonly applied to identify and annotate the genome into chromatin states based on combinatorial patterns of multiple chromatin modifications, in this study, the genome is annotated based on spatiotemporal-combinatorial mCG and/or chromatin-interaction patterns across developing cell types. We first trained an 80-state ChromHMM model using methylation features at a 200 bp resolution from 180 pseudobulk mCG profiles and annotated each genomic bin with one of the states (**Figure 7A**). We investigated the relationship between these states and an existing chromatin state annotation of the genome^42^. Specifically, for each state, we conducted an enrichment analysis for 100-state universal ChromHMM state annotations, which correspond to patterns of histone modifications and variants and DNase I accessibility in diverse cell and tissue types (**Figure 7B**)^42^. Lastly, we identified putative TFs associated with each methylation-based state based on the enrichment of ChIP-seq binding sites for data curated and processed by ChIP-Atlas (**Figure 7C**)^43^. The methylation-based states captured combinations of developmental-stage-specific and cell-type-specific methylation patterns. For example, states 11-18 showed broad demethylation in developing glial cell types and mid-gestational neuronal progenitors or immature neurons. These regions universally gain mCG during the maturation of all neuronal types (**Figure 7A**). Such a pattern is partially driven by the developmental crosstalk between polycomb repression and *de novo* DNA methylation^44^. States 11-18 are enriched in bivalent promoters and polycomb-repressed universal chromatin states (**Figure 7B**), while also enriched in the binding sites of SWI/SNF chromatin remodellers SMARCA4 and SMARCB1, which antagonize the Polycomb Repressive Complex (PRC)^45^. States 11-18 are also significantly overlapped with the EnhA17-19 universal chromatin states that indicate early developmental enhancers (**Figure 7B**). Consistently, these states are enriched for binding sites of early neuronal lineage-specific TFs, including SOX2 and ASCL1 (**Figure 7C**). Together, the results suggest that the dynamics in states 11-18 reflect a transition from polycomb to methylated-mediated repression in some of the regions^44,46^, in addition to the repression and *de novo* methylation at early developmental enhancers.

**Figure 7.**
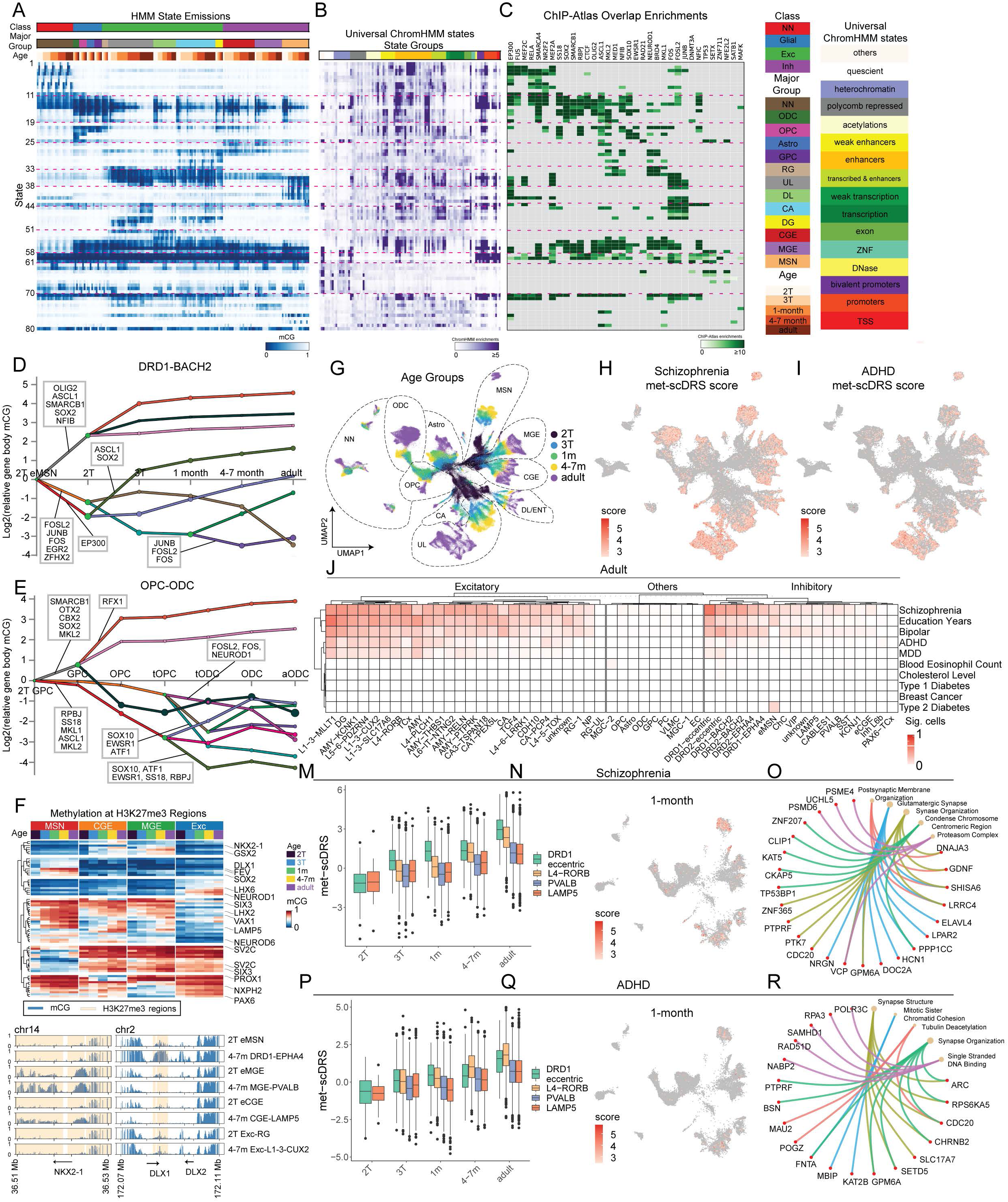
Regulatory landscape of brain development and disease association. (A) ChromHMM-based annotation of the human genome into 80 states (at a 200 bp resolution) based on the dynamic mCG pattern during human brain development. (B-C) Enrichment of universal ChromHMM states (B) and TF binding sites (C). For each state we selected the top-3 most enriched ChIP-Atlas experiments, then computed the median enrichment for each TF across the select experiments. (D-E) DREM reconstruction of transcriptional regulatory dynamics across the differentiation of striosome DRD1-expressing MSNs (D) and across the differentiation of oligodendrocytes in infant brains (E). (F) Trajectory-specific mCG level at H3K27me3 modified regions. (G-I) UMAP dimensionality reduction of the snm3C-seq dataset labeled with age groups and major cell type groups. (J) Fraction of single cells showing significant met-scDRS score for selected neuropsychiatric and non-brain traits in the adult brain. (M-N) Enriched met-scDRS score for SCZ in DRD1-expressing eccentric MSNs in the 1-month-old brain. (O) Selected genes whose gene body mCG level most strongly correlated with met-scDRS scores for SCZ in the 1-month-old brain. (P-Q) Enriched met-scDRS score for ADHD in DRD1-expressing eccentric MSNs in the 1-month-old brain. (R) Selected genes whose gene body mCG level most strongly correlated with met-scDRS scores for ADHD in the 1-month-old brain.

States 33-51 are associated with developmental loss of mCG (compared to the 2T stage) in combinations of neuronal cell types, with states 33 and 44-51 demethylated in excitatory neurons, states 38-43 demethylated specifically in MSNs, states 35-37 demethylated in both excitatory and inhibitory neurons (**Figure 7A**). The majority of these states are enriched in enhancer-associated universal chromatin states (**Figure 7B**), and the binding sites of activity-dependent TFs FOS, FOSL2, and JUNB, again highlighting the critical role of AP-1 TFs, potentially driven by neuronal activity, in shaping the neuronal regulatory landscape (**Figure 7C**).

We also explored the joint ChromHMM modeling of mCG and total chromatin contact at 5 kb resolution, and found that mCG generally shows greater cell-type and developmental dynamics than chromatin contacts (**Figure S7A-C**). Key developmental patterns were recapitulated using this multi-modal modeling approach. For example, the developmental gain of mCG at bivalent chromatin was captured in joint states 55-63 (**Figure S7A**); the neuronal-specific developmental demethylation was detected by states 65-71. Together, the application of ChromHMM and ChIP-Atlas across diverse cell types identified crosstalks mediated by transcriptional cofactors, such as PRC and SWI/SNF complexes, a type of regulatory dynamics that can be overlooked by TF binding motif analysis.

### Regulatory dynamics of neural differentiation

We used DREM to infer a dynamic regulatory map of mCG changes for representative differentiation trajectories by jointly modeling mCG dynamics at DMRs and TF–DMR associations (determined by ChIP-Atlas^43^) as a sequence of bifurcation events regulated by TFs^26^. In line with the finding from ChromHMM analysis, across multiple trajectories (**Figure 7D-E and S7D-F**), regions gaining mCG in mid-gestational development are enriched for binding motifs for the Polycomb repressive complex 2 (PRC2) antagonists, SMARCB1 and SMARCA4. This suggests that PRC2 is developmentally reduced in these regions. Regions gaining mCG in mid-gestation are also enriched in the binding sites of early patterning TFs such as ASCL1 and SOX2. In neuronal trajectories, the binding sites of pro-glial factors OLIG2 and NFIB also gain mCG (**Figure 7D and S7D-F**). The DREM analysis recapitulated the finding that the binding sites of neuronal activity-dependent TFs, including AP-1 factors FOSL2 (Score: 7.7×10^-15^), JUNB (Score: 5.8×10^-12^), FOS (Score: 4.4×10^-10^), are more strongly enriched in the demethylated regions in the differentiation of DRD1-Striosome than in the differentiation of DRD1-Matrix (FOSL2 Score: 6.1×10^-5^, FOS Score: 3×10^-4^, JUNB Score: 0.014) (**Figure 7D and S7D**). In regions demethylated during differentiation, DREM identified enriched binding of lineage-specific TFs, such as SOX10 for OPC/ODC, and MEF2C for MGE-derived inhibitory neuron trajectories (**Figure 7E and S7F**).

### Trajectory-specific methylation dynamics at Polycomb repressed regions

During development, DNA methylation and PRC2-mediated gene silencing both contribute to dynamic regulation of gene expression. To identify the relationship between these two major epigenetic modes of dynamic repression, we examined cell-type-specific mCG levels at genomic regions enriched in PRC2-associated histone mark H3K27me3, in DLPFC. We focused on promoter-proximal regions, within 2 kb of annotated transcription start sites (TSS)^47^. We identified 70 TSS-overlapping regions showing strong cell-type specificity, defined by a developmental-averaged mCG difference greater than 0.2 (**Figure 7F and Table S4**). At some lineage-specific genes, such as NKX2-1, we observed a paradoxical pattern of hyper-mCG in eMGE and MGE-PVALB, both derived from MGE progenitors that express a high level of NKX2-1. The hyper-mCG at NKX2-1/*Nkx2-1* in PV+ neurons is observed in both human and mouse^48^. This pattern is consistent with early expression of NKX2-1/*Nkx2-1* in MGE progenitor cells prior to our earliest samples, and reflects the antagonism of PRC2 repression by *de novo* methylation. However, not all lineage-specific methylation patterns can be explained by past expression in progenitors. For example, DLX1 shows a specific gain of mCG in MSN neurons, yet DLX1 is broadly expressed in GEs^49^. This suggests that transitions from PRC2-associated repression to DNA methylation–associated repression may occur even in the absence of strongly cell-type-restricted expression at the corresponding developmental stage.

### Neuropsychiatric risk across single cells in developing brains

We reasoned that our high-resolution data about developmentally dynamic, cell type-specific DMRs could help to interpret the functional significance of genetic loci associated with risk for neuropsychiatric traits. We employed a recently developed gene set-based, single-cell resolution method, met-scDRS (methylation single cell Disease Relevance Score) (**Figure 7G-I**)^27^. Met-scDRS does not depend on pseudobulk summaries of single cell data, in contrast to other methods that test for polygenic heritability across gene sets, such as MAGMA^50^. The specificity of the met-scDRS is supported by strong enrichment signals at risk loci for neuropsychiatric traits, and then absence of enrichment for non-brain-related traits (**Fig. 7J**). In our previous study, by applying stratified linkage disequilibrium score regression to DMRs and chromatin loop-connected DMRs^51^, we found that active regulatory regions in post-mitotic neurons show a greater enrichment of the genetic risk of schizophrenia (SCZ) and bipolar disorder (BIP) than in neural progenitor cells^15^. We recapitulated this observation using gene set-based met-scDRS analysis, showing a general pattern that the risk of SCZ and BIP, in addition to ADHD (Attention-Deficit Hyperactivity Disorder) and MDD (Major Depressive Disorder) increases in neuronal but not non-neuronal cells, between mid-gestation and adulthood (**Figure S7G-J**). The visualization of met-scDRS scores using single-cell embedding also highlighted the increase in met-scDRS scores along with cellular maturation (**Figure 7G-I**).

The clustering of diseases by the pattern of met-scDRS scores revealed two classes of neuropsychiatric disorders. Risk loci of SCZ and BIP were enriched in both excitatory and inhibitory neurons (**Figure 7J, S6K-L**). By contrast, risk loci for ADHD and MDD (Major Depressive Disorder) were primarily enriched in excitatory neurons (**Figure 7J, S7K-L**). Among inhibitory neurons in the adult brain, DRD1-eccentric and DRD2-Striosome neurons are among the most strongly associated with SCZ, with 67.5% and 44.9% of cells significantly associated with SCZ, respectively (**Figure 7J**). Notably, in all age groups, the LGE-derived MSNs are more strongly enriched in the risk of SCZ than CGE and MGE-derived inhibitory neurons (**Figure 7J** and **S7K-L**) (p-value < 2.2e-16, chi-square test). The met-scDRS results were also supported by LDSC partitioned heritability analysis using loop-connected DMRs identified from pseudotime bulk profiles (**Figure S7M-N**)^51,52^.

We showcased the met-scDRS pattern of SCZ and ADHD across 1-month old brain cell populations, highlighting the strong enrichment of SCZ polygenic risk in MSNs, especially the DRD1-eccentric population (**Figure 7M-O**). Although MSNs in 1-month old brains are also relatively enriched in ADHD heritability, the met-scDRS scores of ADHD in excitatory neurons generally exceed those of MSNs in 1-month and adult brains (**Figure 7P-R**). To identify genes that drive the disease risk profile in the 1-month old brain, we extracted top genes whose gene body mCG level most strongly correlated with met-scDRS scores for SCZ and ADHD across cell populations and determined the gene ontologies associated with these genes (**Figure 7O, R**). We found that genes related to synapse machinery, such as HCN1 and NRG,N are among the genes most correlated with met-scDRS scores and were also identified as SCZ risk genes by GWAS (**Figure 7O**)^53^. Genes whose gene body mCG level most strongly correlated with met-scDRS for ADHD include transcriptional and chromatin regulators implicated in modulating neuronal functions, including RPS6KA5, KAT2B, SETD5, and POGZ (**Figure 7R**). In summary, using the novel gene set-based met-scDRS approach, our analysis recapitulated the increase of polygenic risk of SCZ, BIP, ADHD, and MDD across development, highlighted the strong enrichment of disease risk in MSNs, and identified genes with strong contributions to scDRS scores.

## Discussion

The single-cell and spatial 3D-epigenomic dataset generated from GEs, developing basal ganglia and inhibitory neurons, fills a critical gap in spatiotemporal data resources for the study of gene regulation during human brain development. The 3D-genome conformation of multiple brain regions in developing human brains has been studied using bulk Hi-C profiling^35,54^. However, single-cell studies of the 3D-genome conformation have so far been applied to developing human DFC, HIP^15^, and postnatal cerebellum^16^, and have not sampled the progenitor zones of inhibitory neurons. In addition, sampling only DFC precludes the study of regulatory signatures of cortical arealization. Our sampling of GEs and excitatory VZ extended the analysis of the developmental trajectory of both excitatory and inhibitory neurons. In addition, sampling of the key 1-month postnatal period revealed novel developmental dynamics, including transient 3D genomic remodeling in a subset of neuronal types and during the transition from OPC to ODC, both of which were enriched in 1-month-old brain samples. Our work highlights the dynamic nature of the neonatal and infant brain across all cell types. Lastly, using the 3D-epigenomic dataset, our study generated comprehensive functional annotations, including DMRs, chromatin loops, and chromatin domain boundaries, which can serve as valuable resources for dissecting gene regulation and identifying genetic variants associated with neurodevelopmental and neurological diseases.

### 3D-epigenomic landscape of developing MSN subtypes

Recent brain cell atlasing using cutting-edge single-cell technologies have identified new cell types (e.g., eccentric MSN) in the basal ganglia and determined the molecular properties of cells associated with striatal structures (e.g., striosome vs. matrix)^55–57^. Our study expanded such characterizations to 3D-genome and methylome modalities throughout the basal ganglia development. Despite the traditional classification of MSNs into D1 and D2 types based on the expression of the dopamine receptor genes DRD1 and DRD2, our unbiased classification using either methylome or 3D-genome information reproducibly identified MSNs in striosomal and matrix compartments exhibiting greater epigenomic differences, independent of their D1/D2 types. Meanwhile, eccentric MSNs are a clear outlier, distinct from all other MSN subtypes. The analysis of developmental DMRs revealed highly distinct TF regulatory programs during the differentiation of MSN subtypes, with eccentric MSNs showing the greatest similarity to immature MSNs; striosomal and matrix MSNs show drastic differences in the timing of the emergence of neuronal activity-dependent AP-1 TF binding signatures. Interestingly, eccentric MSNs are strongly enriched for polygenic risk for SCZ and BIP, suggesting that this less abundant MSN subtype should be further investigated for its role in neurodevelopmental disorders.

### Synchronized maturation of MSNs

Multiple analyses in this study have found that MSNs exhibit unusual homogeneity within an age group and a discrete separation between age groups, which are clearly distinct from those of all other types of excitatory and inhibitory neurons analyzed so far. While developing inhibitory interneurons traverse large 3D-genome pseudotime distances in mid-gestation, and traverse large methylome pseudotime distances in late-gestational to early infant development, such an intra-donor heterogeneity was conspicuously absent for MSNs. The results indicate that MSN development proceeds in a highly synchronous manner, leading to reduced developmental heterogeneity at any given age compared with other cell types. This synchronization likely reflects the need for early, precise establishment of basal ganglia circuitry. Therefore, the developmental trajectory of MSNs contrasts with existing studies of human brain development, which consistently find heterogeneity in differentiation and maturation states among neurons sampled from the same donor.

### Transient 3D genome reorganization

Several recent studies have found that the enrichment of short-range or long-range chromatin interactions (i.e., SE or LE, respectively) is a major signature distinguishing the chromatin organization of neuronal and non-neuronal cells^15,16,19^. While all previously known SE to LE changes (or *vice versa*) are monotonic and are associated with terminal cell-type differentiation (e.g., to post-mitotic neurons or astrocytes) or aging, this study has uncovered pronounced heterogeneity in SE/LE states in glial cells, and a transient switch to SE during the transition between OPC and ODC. Motivated by our previous findings that SE is associated with increased numbers of chromatin loops and domain boundaries, we speculated that the transient conversion to SE could increase cellular transcriptomic output to support the differentiation from OPC to ODC. Such a speculation was consistent with the enrichment of SE in neuronal cell types and the observation that neuronal nuclei are transcriptionally more active than glial nuclei.

### The role of 3D genome dynamics in brain development

Our study has described intriguing correlations between brain cellular dynamics, global 3D genome conformation, nuclear volume, and transcriptomic output. Mechanistic studies will be necessary to distinguish causation from correlation and to determine the role of nuclear and 3D genomic remodeling in brain development. Open questions include the mechanisms that control the switching between SE and LE states. Although the SE state likely results from an increased loading of the cohesin complex onto the chromatin^58^, the developmental signals that instruct the process and the mechanistic steps of cohesin dynamics are unknown. Furthermore, it is unknown whether any causal relation exists between SE conformation, genome-wide transcriptional activation, and nuclear volume, despite the fact that strong correlations have been observed in pairs of observations in various contexts^38,59^. Lastly, further works are needed to identify how neural lineage-specific TFs instruct the remodeling of global chromatin conformation and nuclear morphology, and whether such structural reconfigurations are essential for the differentiation and maturation of neuronal and glial cell types.

## Acknowledgement

We thank Kenneth X. Probst for illustration. This work is supported by the National Institute of Mental Health (NIMH) awards U01MH130995 to C.L. M.F.P, J.E., and E.A.M.; R01MH125252 to C.L. and B.P.; RF1MH130461 to C.L; NIH Office of the Director DP5-OD031878 to B.B., Chan Zuckerberg Initiative AICP-0000000175 to B.B. and Q.Z. The Flow Cytometry Core Resource is supported by the Eli and Edythe Broad Center of Regenerative Medicine and Stem Cell Research Center (BSCRC) at the University of California Los Angeles. Confocal microscopy images were acquired at the UCSF Innovation Core at the Weill Institute for Neurosciences. This publication was supported by and coordinated through the Brain Initiative Cell Atlas Network (BICAN).

## Data and code availability

Fastq files for the snm3C-seq data, processed single-cell methylation profiles, and processed single-cell chromatin conformation profiles are available through the NeMO archive under accession nemo:dat-n5emc24 (https://assets.nemoarchive.org/dat-n5emc24). DNA methylation and chromatin conformation profiles aggregated for annotated cell types are available through NCBI GEO accession. The data are available for visualization at https://brainome.ucsd.edu/bg-epigenome-development-atlas. An interactive browser for visualizing the ChromHMM analysis results is available at https://github.com/luke0321li/BICAN_ChromHMM. MERFISH and chromatin tracing data will be available on Dryad at https://doi.org/10.5061/dryad.3tx95x6wd and can be accessed by reviewers at http://datadryad.org/share/LINK_NOT_FOR_PUBLICATION/B0Eg_TOkYfJMyDacVNU QRydDXOhzM6njs9sHFNAwWA0

## Supplemental Figures and Tables

**Figure S1.**
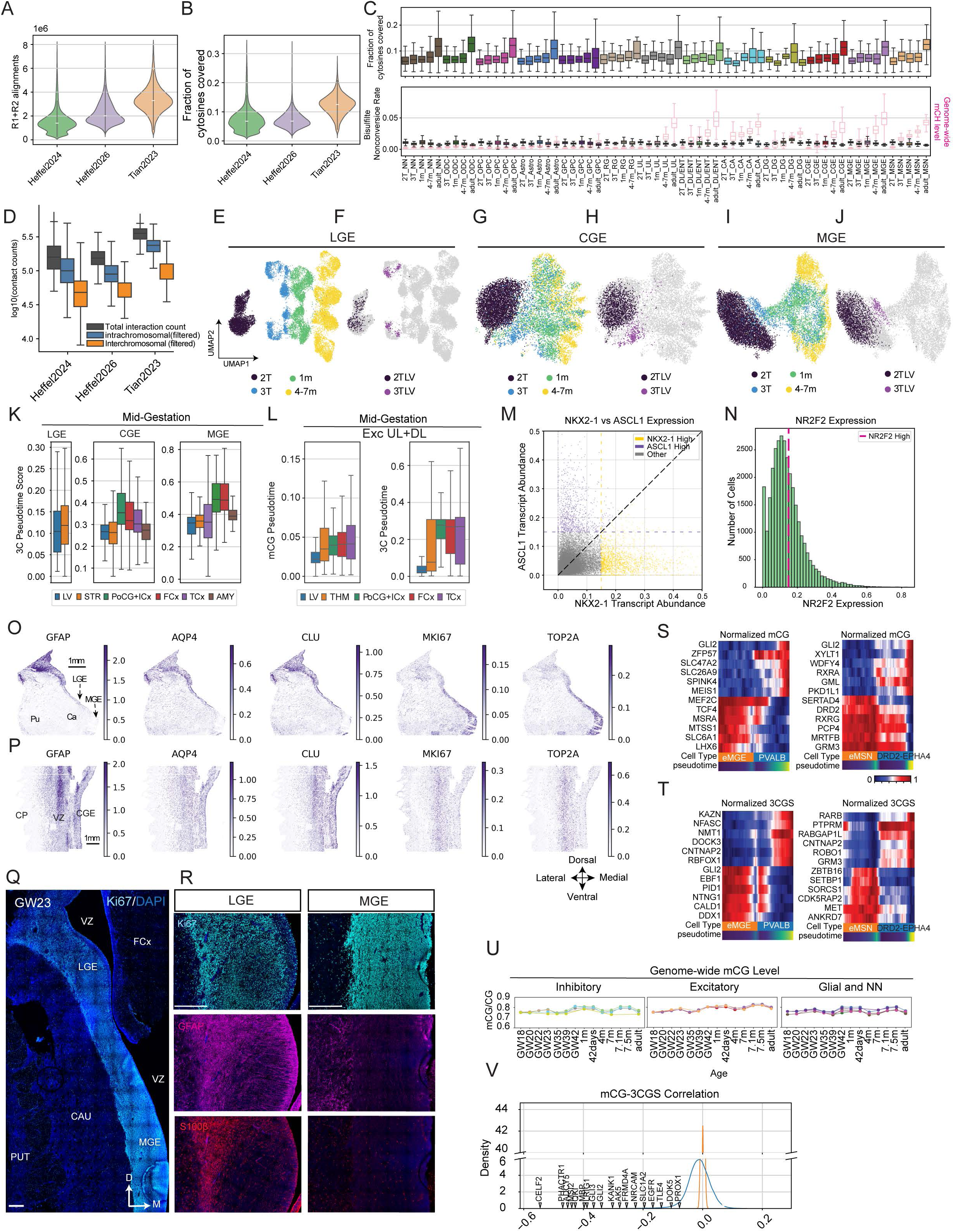
Characterization of the developing basal ganglia and inhibitory neurons using snm3C-seq and spatial transcriptomics, related to Figure 1. (A) The number of uniquely mapped methylome reads in each dataset integrated in this study. (B) The fraction of cytosines covered by cytosine positions in each dataset integrated in this study. (C) Coverage (top), bisulfite non-conversion rate (bottom), and mCH level (bottom) for major cell groups for each age group. (D) The number of total, intrachromosomal, and interchromosomal chromatin contacts in each dataset integrated in this study. (E-F) UMAP dimensionality reduction of LGE-derived cells labeled for age groups (E) or cells dissected from LV (F). (G-H) UMAP dimensionality reduction of CGE-derived cells labeled for age groups (G) or cells dissected from LV (H). (I-J) UMAP dimensionality reduction of MGE-derived cells labeled for age groups (I) or cells dissected from LV (J). (K) Distributions of single-cell 3C pseudotime scores for GE-derived neurons across brain regions. (L) Distributions of single-cell mCG (left) and 3C (right) pseudotime scores for excitatory neurons across cortical areas. (M) Cells with high NKX2-1 expression were selected by requiring > 0.15 normalized counts and greater than the abundance of ASCL1 transcripts. Cells with high ASCL1 expression were selected by requiring > 0.15 normalized counts and greater than the abundance of the NKX2-1 transcript. (N) Cells with high NR2F2 expression were selected by requiring > 0.15 normalized counts. (O-P) Transcript abundance of astrocyte and proliferation markers on the anterior section (O) and the posterior section (P). (S) Normalized mCG for marker genes that distinguish eMGE and MGE-PVALB (left), or eMSN and DRD2-Matrix (right). Normalized 3CGS for marker genes that distinguish eMGE and MGE-PVALB (left), or eMSN and DRD2-Matrix (right). (Q-R) An anterior section of a 23 GW section was stained for a proliferation marker Ki67 (Q-R), GFAP (R), and S100B during the differentiation and maturation of inhibitory (left), excitatory (mid), and glial cells (right). (V) The correlation between gene body mCG and 3C Gene Score (3CGS) highlights the strong inverse correlation at cell-type marker genes.

**Figure S2.**
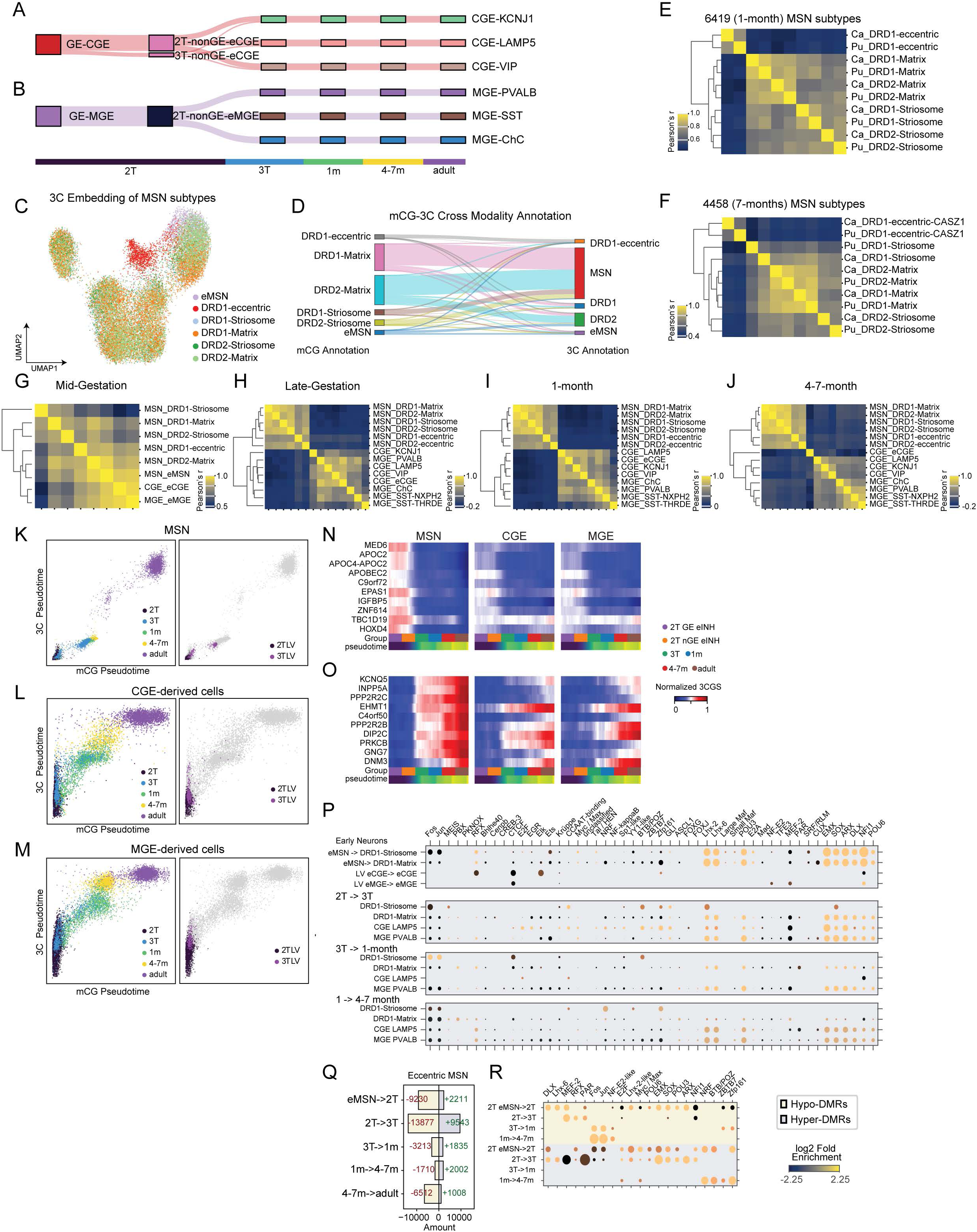
Comparison of development dynamics across MSN subtypes and interneurons. Related to Figure 2. (A-B) The specification of CGE (A) and MGE (B) subtypes in late-gestation. (C) UMAP embedding of MSN subtypes using 3C information. (D) mCG-3C cross-modality comparison of MSN neuron classifications. (E) Correlation matrix of MSN subtypes identified in an 1-month old donor (6419), computed using scaled gene body mCG. (F) Correlation matrix of MSN subtypes identified in a 7-month old donor (4458), computed using scaled gene body mCG. (G-J) Correlation matrices of inhibitory neuron subtypes identified in mid-gestational (G), late-gestational (H), 1-month-old (I), and 4–7-months-old (J) donors. (K-M) Comparison of pseudotime scores computed using mCG and 3C modalities across the differentiation of LGE-derived MSNs, and CGE- and MGE- derived interneurons, including cells derived from adult donors. (N-O) Scaled 3CGS of selected marker genes during the differentiation of LGE, CGE, and MGE-derived cell populations. The marker genes were selected for loss (N) and gain of 3CGS (O) during MSN differentiation. (P) TF binding motif enrichments in trajectory hyper-DMRs. (Q-R) Trajectory DMRs and associated TF enrichment (R) identified across the differentiation of eccentric MSNs.

**Figure S3.**
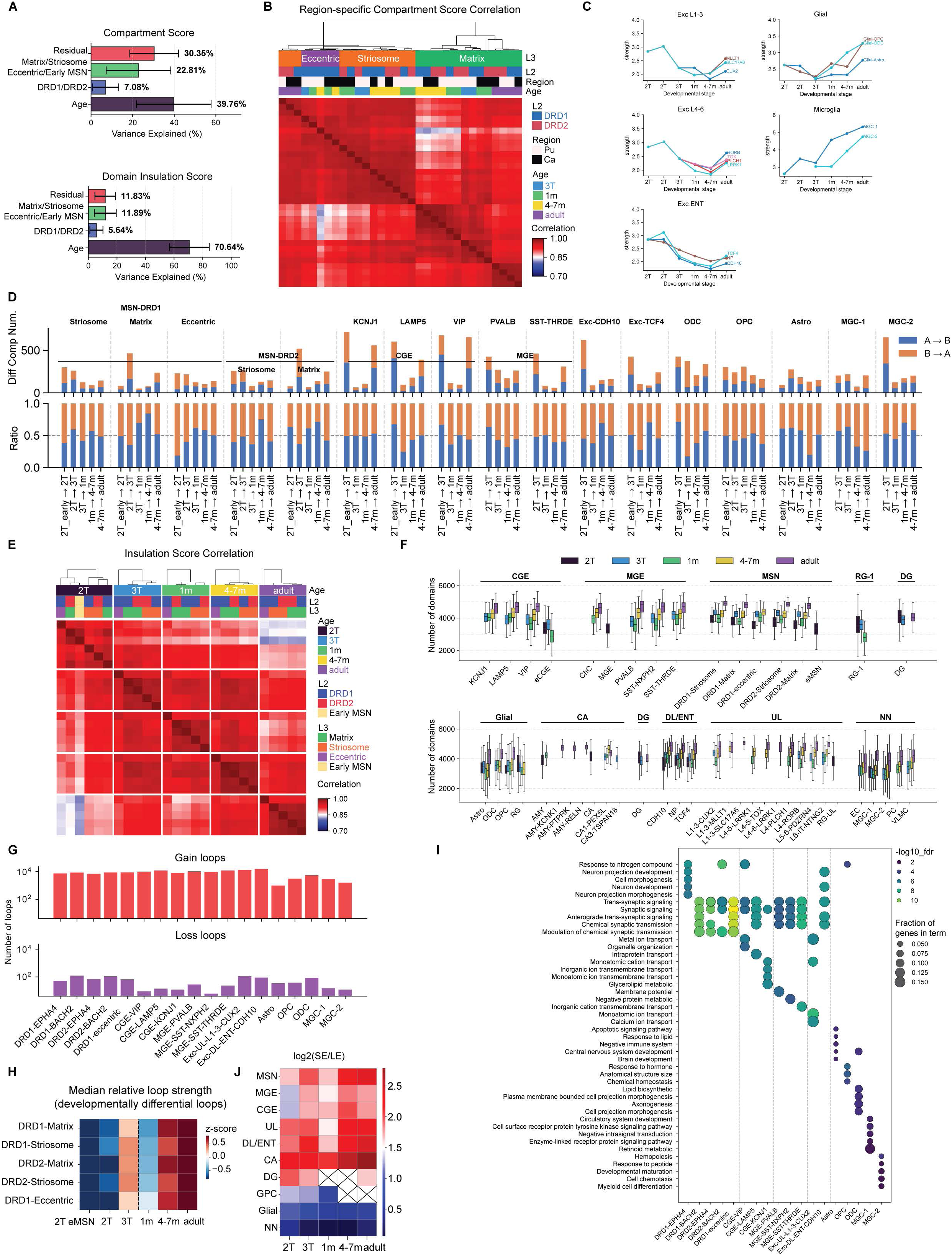
Chromatin changes across developmental stages. Related to Figure 3. (A) Variance partitioning based on ANOVA, showing the relative contributions of age, DRD identity, and MSN subtype to variation in 100-kb compartment scores and 25-kb insulation scores. (B) Genome-wide Pearson correlations of A/B compartment scores across MSN subtypes, stratified by Pu and Ca regions. (C) Developmental changes in compartment strength across excitatory neuronal lineages, glia, and microglia. (D) Number of differential compartments and the ratio of B-to-A and A-to-B compartment switches across cell types. (E) Genome-wide Pearson correlations of 25-kb bin insulation scores across MSN subtypes. (F) Number of domains identified across cell types and developmental stages. (G) Number of differential chromatin loops (gains and losses) across major developmental lineages. (H) Relative strength of developmentally regulated loops across MSN subtypes. (I) Gene Ontology (GO) enrichment analysis of genes associated with loop gains within ±2 kb of promoters. (J) Average log₂ ratio of short- to long-range interactions across cell types and developmental stages.

**Figure S4.**
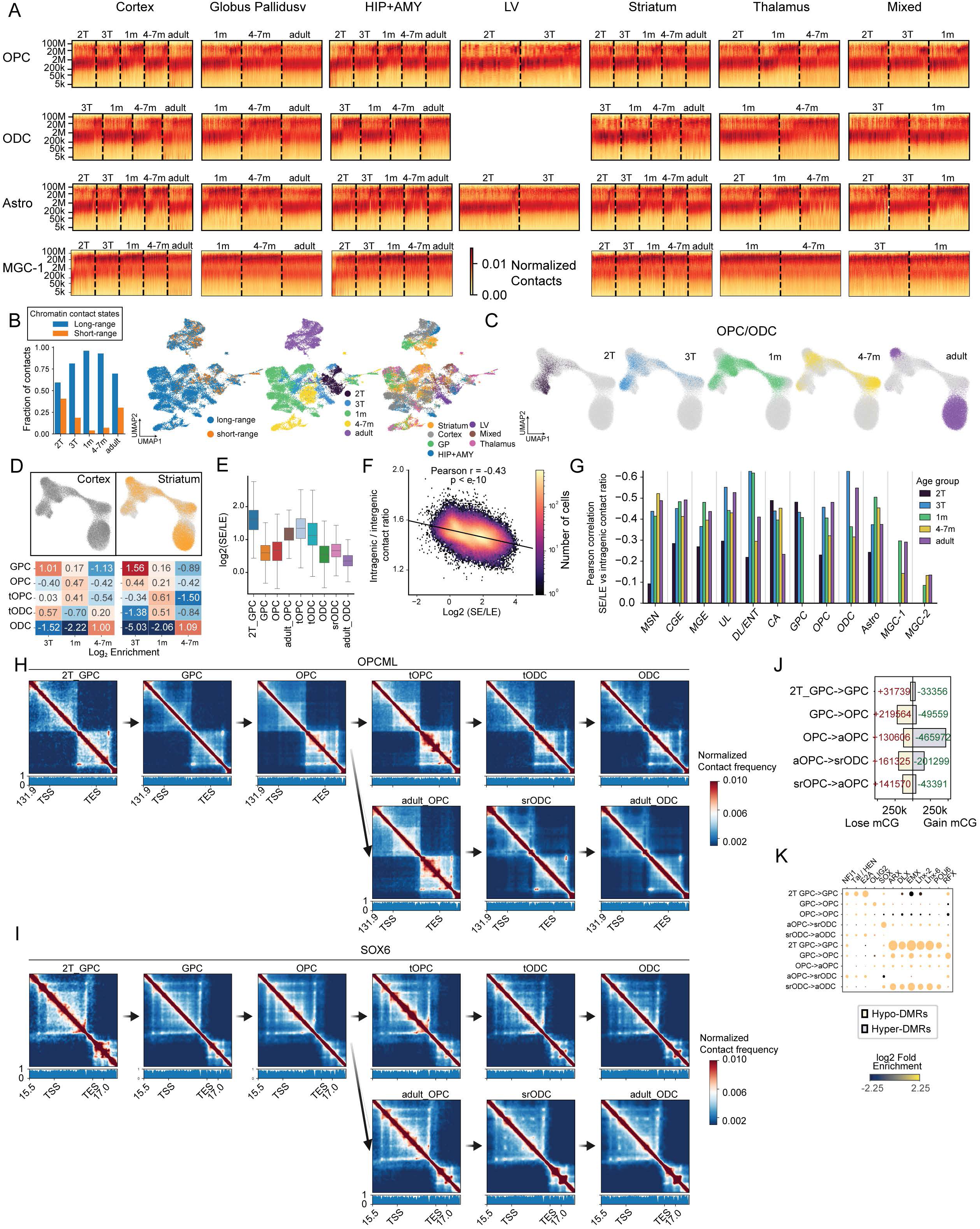
Epigenomic changes in OPC/ODC during development. Related to Figure 4. **(A)** Heatmaps depict contact frequency by genomic distance in individual cells across developmental stages and brain regions for OPCs, ODCs, astrocytes, and microglia-1. **(B)** Proportion of short- and long-range chromatin interactions (SE/LE) in astrocytes across developmental stages. **(C)** UMAP embedding of the 3C modality across developmental stages for OPC and ODC cells. **(D)** Log₂ enrichment of OPC/ODC subtypes across developmental stages in cortex and striatum. Enrichment was calculated as the ratio of the proportion of each cell type at a given developmental stage to its overall frequency across all stages. **(E)** log₂(SE/LE) across OPC and ODC subtypes. **(F-G)** Correlation between the log₂(SE/LE) and the intragenic/intergenic contact ratio across all cells **(F)** and within major cell types across developmental stages **(G)**. **(H-I)** Local chromatin confirmation dynamics at OPCML and SOX6 during the differentiation of ODC in infant and adult brains. **(J-K)** Trajectory-DMRs **(J)** and TF binding motifs **(K)** identified across the differentiation of ODC in adult brains.

**Figure S5.**
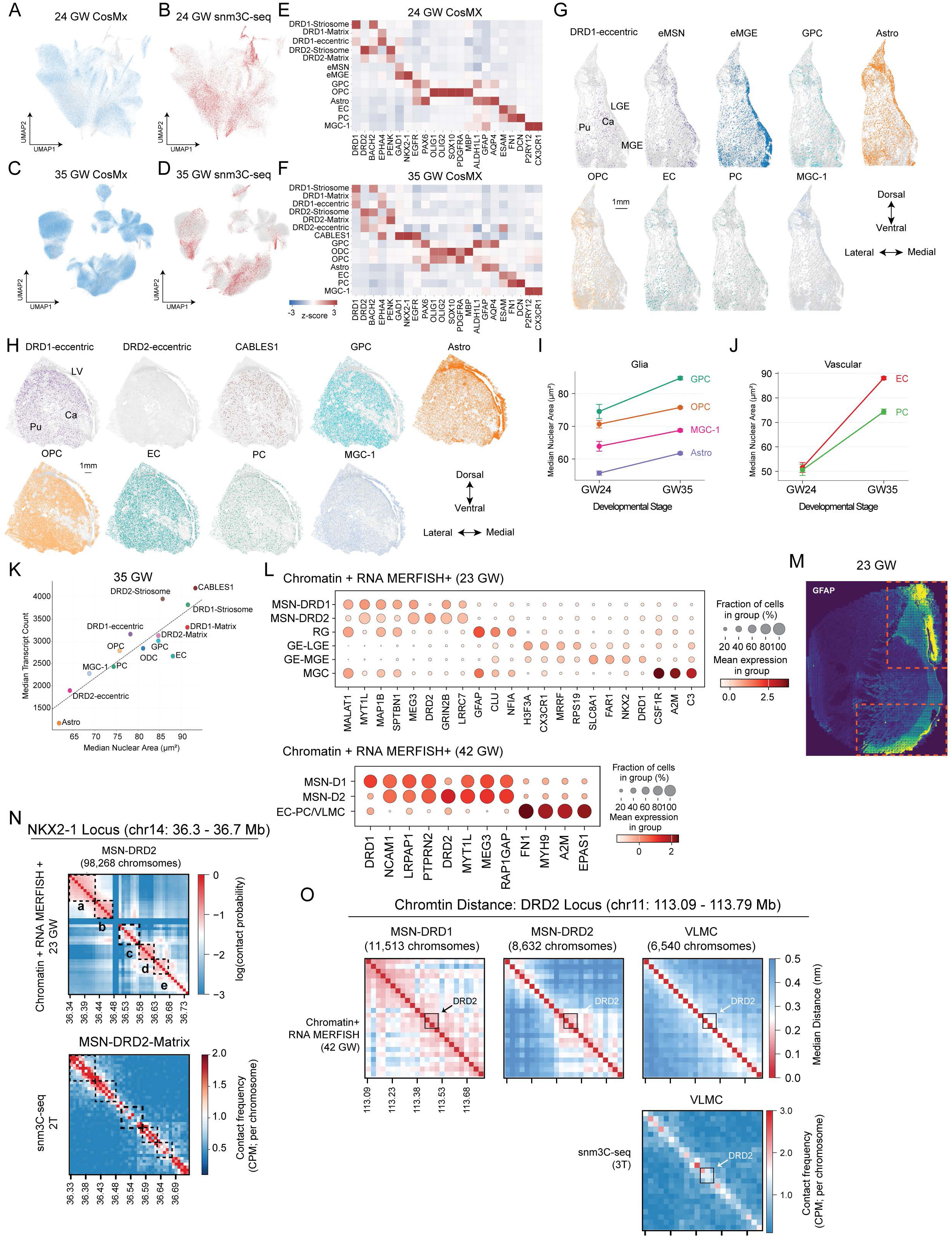
Characterize the developing basal ganglia using highly multiplexed spatial transcriptomic and chromatin+RNA MERFISH. Related to Figure 5. (A–D) UMAPs showing the integration of snm3C-seq cells with CosMx spatial transcriptomic profiles in the 24 GW and 35 GW samples. (E–F) Z-score expression values of marker genes used to identify major cell types in the integrated dataset. (G–H) Spatial maps of annotated cell populations in the 24 GW and 35 GW samples. (I–J) Nuclear area distributions for glial and other non-neuronal cell types across developmental stages. (K) Relationship between nuclear area and transcriptional activity in the 35 GW sample. (L) Marker-gene expression supporting major cell-type assignments in the combined chromatin tracing + MERFISH+ experiments. (M) Spatial map of GFAP expression in the 23 GW sample. (N) Chromatin tracing contact probability at the NKX2-1 locus across cell types in the 23 GW sample, alongside the corresponding snm3C-seq contact matrix. (O) Median chromatin distance from chromatin tracing at the DRD2 locus in the 42 GW sample, together with 3C contact maps for VLMC in the third trimester.

**Figure S6.**
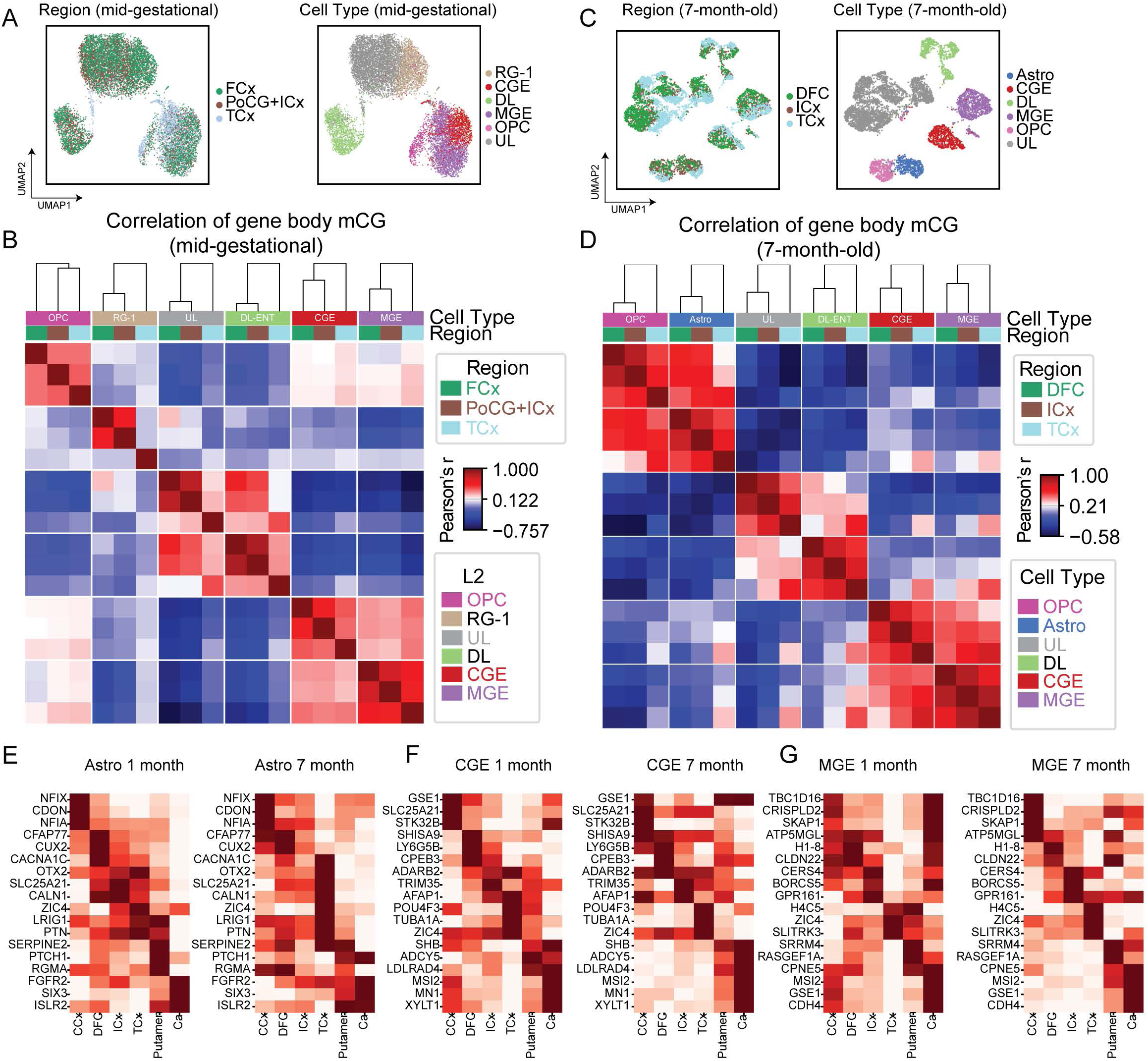
Epigenomic signatures of cortical arealization in developing human brains. Related to Figure 6. (A) mCG UMAP embedding of 23 GW old donor 2301 for major cell groups represented in the cortical regions colored by region (left) and lineage (right). (B) Pearson correlation of DFC, PoCG+ICx, and TCx across all major cell groups represented in these three dissections. (C) mCG UMAP embedding of 7 month old donor 4285 for major cell groups represented in the cortical regions, colored by region (left) and lineage (right). (D) Pearson correlation of DFC, ICx, and TCx across all L2 lineages represented in these three regions. (E-G) Top mCG genes shared by 1-month and 7-month samples for astrocyte (E), CGE-derived neurons (F), and MGE-derived neurons (G) regional specificity.

**Figure S7.**
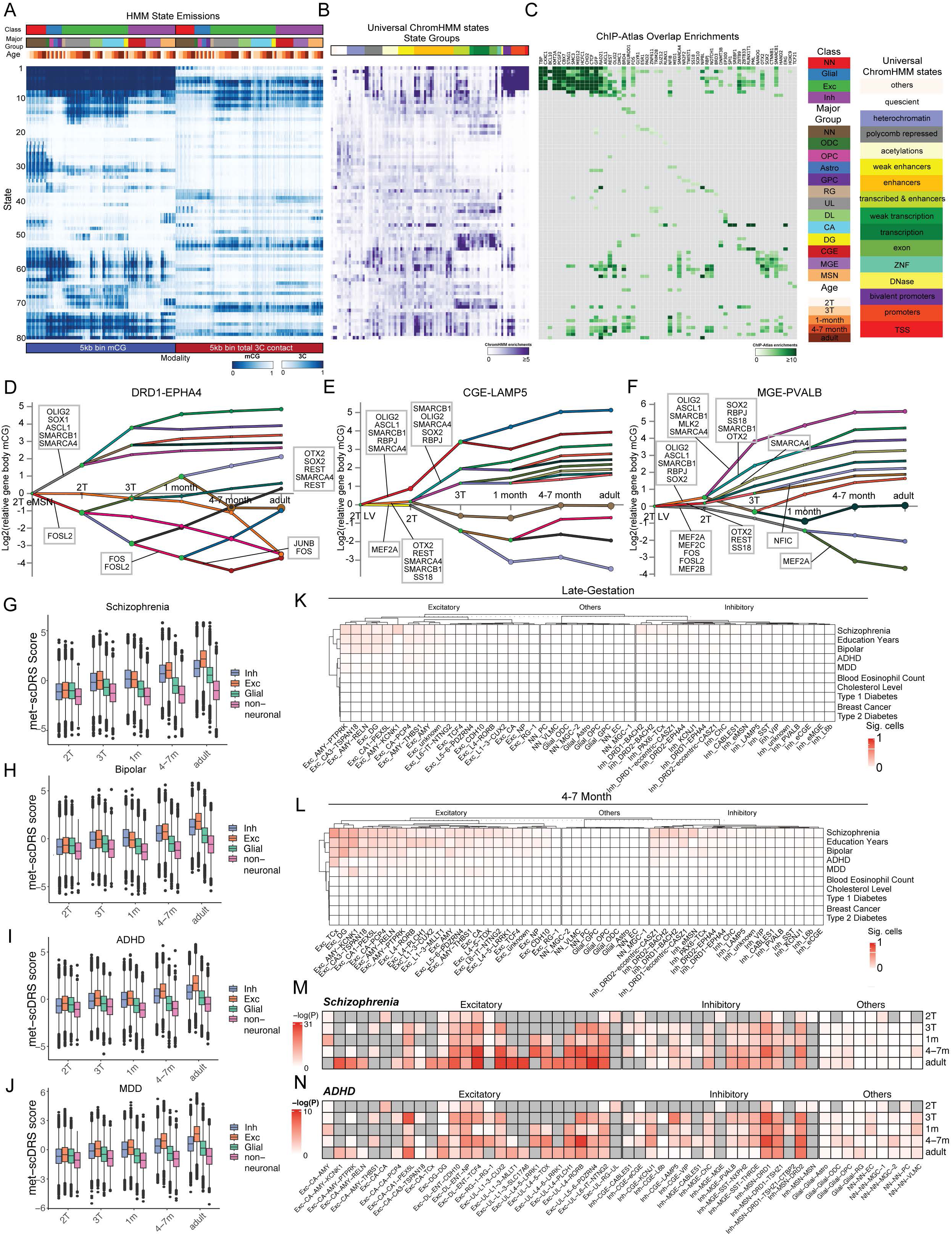
Regulatory landscapes of developing basal ganglia and inhibitory neurons. Related to Figure 7. (A) ChromHMM-based annotation of the human genome into 80 states (at a 5kbp resolution) based on the joint dynamic mCG and 3C patterns during human brain development. (B-C) Enrichment of universal ChromHMM states (B) and TF binding sites (C). For each state we selected the top-3 most enriched ChIP-Atlas experiments, then computed the median enrichment for each TF across the select experiments. (D-F) DREM reconstruction of transcriptional regulatory dynamics across the differentiation of matrix DRD1-expressing MSNs (D), CGE-LAMP5 interneurons (E), and MGE-PVALB interneurons (F). (G-J) Quantification of met-scDRS score in inhibitory neurons, excitatory neurons, glial cells, and non-neural cells, for SCZ (G), BIP (H), ADHD (I), and MDD (J). (K-L) Fraction of single cells showing significant met-scDRS score for selected neuropsychiatric and non-brain traits in the late-gestational (K) and 4-to-7 month old (L) human brains. (M-N) Enrichment of polygenic risks for SCZ (M) and ADHD (N) in pseudobulk methylome profiles determined by the LDSC partitioned heritability analysis.

**Table S1.** Human Brain Specimen used in this study.

**Table S2.** Metadata of the integrated snm3C-seq dataset.

**Table S3.** Cortical-regional-specific methylation signatures.

**Table S4.** Cell-type-specific DNA methylation in polycomb-regulated regions.

