## Supplementary material for "A Single-Cell and Spatial 3D Multi-omic Atlas of Developing Human Basal Ganglia and Inhibitory Neurons": Material & Methods

#### Tissue Collection

Human brain tissue was obtained with post-mortem intervals <24 h through the University of California San Francisco (UCSF; second-trimester samples) and the NIH NeuroBioBank (third-trimester and infant samples). Written informed consent was obtained in accordance with institutional regulations. All procedures were reviewed by the UCSF Committee on Human Research and approved by the Human Gamete, Embryo and Stem Cell Research Committee (IRB GESCR #10-02693) and related UCSF IRBs (#20-31968). For prenatal donations, consent for the clinical procedure was obtained prior to tissue donation consent. Donors were informed that participation would not affect clinical care and that neither donors nor clinical teams would receive compensation or benefits. UCSF samples were evaluated by a neuropathologist and classified as gross anatomical control. Additional specimens were accessed through the NIH NeuroBioBank under their institutional approvals.

For snm3C-seq3 and MERFISH, dissected tissue blocks were flash-frozen in liquid nitrogen and stored at −80 °C. Tissue was cryosectioned coronally at −20 °C, and regions of interest were microdissected using a surgical microscalpel. Approximately 30–50 mg of tissue were collected per region into RNAse-free 1.5 mL tubes and immediately stored at −80 °C until processing. For CosMx analyses, fixed-frozen tissue was used. Tissue blocks were fixed in 4% paraformaldehyde (PFA) for 48 h, cryoprotected in 30% sucrose, embedded in OCT, and sectioned coronally at 10 μm before mounting on glass slides. For anatomical validation, multiple sections spanning the anterior–posterior axis of each block were cresyl violet–stained. Anatomical landmarks, including the lateral ventricle, ganglionic eminences, and basal ganglia subregions, were identified using the BrainSpan Atlas of the Developing Human Brain and the Bayer & Altman histological atlas series.

#### Single-nucleus methyl3C sequencing

Each snm3C-seq reaction was performed with a brain tissue aliquot with a weight between 30-50 mg using methods described in Heffel et al [1]. All snm3C-seq reactions were carried out with anti-NeuN labelling using anti-NeuN antibody (PE-conjugated, clone A60, Millipore-Sigma number FCMAB317PE). A detailed bench protocol for snm3C-seq is provided through protocol.io (https://doi.org/10.17504/protocols.io.kqdg3x6ezg25/v1). Snm3C-seq libraries were sequenced at the New York Genome Center using the Novaseq X Plus sequencer with 25B 300-cycle reagent kits.

***Single-Cell Spatial Transcriptomics with CosMx Platform***

**CosMx RNA Slide preparation and processing**

Three independent 10µm-thick coronal human brain sections, a more anterior 23 GW, a more posterior 24 GW, and a more anterior 35 GW section, were obtained from cryostat sectioning at -20 °C. Samples were 4% PFA-fixed and mounted onto positively charged Superfrost slides and stored at -80ºC until ready for use. Tissue-mounted slides were processed as described in the "Fresh Frozen" section of the CosMx™ SMI Manual Slide Preparation for RNA Assays manual (Bruker Spatial Biology, MAN-10184-04) using reagents and components provided in the CosMx™ FFPE Slide Prep Kit (RNA) (Bruker Spatial Biology, 121500006). Once ready, slides containing the tissue sections were briefly defrosted at room temperature for 2 minutes and washed thrice in PBS within 50mL Falcon® Tubes. To increase tissue adhesion to slides, slides were baked at 60°C for 40 minutes, and then washed in PBS an additional time. To rehydrate, slides were washed in 50% ethanol for 5 minutes and then stored in 70% ethanol at 4°C overnight. The following day, slides were washed in 100% ethanol twice and then left on a benchtop to air dry for 10 minutes. Antigen retrieval was performed by placing slides within a preheated coplin jar containing NanoString 1X Target Retrieval Buffer, and heating the coplin jar within an Instant Pot pressure cooker at 100°C for 10 minutes. Slides were quickly washed in DEPC water for 5 minutes and then 100% ethanol for 3 minutes. Following a 30-minute air dry step, tissue permeabilization was achieved by following the manual's "Fresh Frozen tissue types" guide, which entailed using a fresh dilution of Proteinase K at 3 μg/mL working concentration. Once prepared, 400μL of this digestion solution was directly applied onto the tissue, and samples were incubated for 30 minutes at room temperature. Following two PBS washes, freshly prepared and vortexed fiducials were applied to the tissue sections for 5 minutes, and tissue sections were protected from light from this point forward. After another PBS wash, a quick fixation was done by keeping the slides in 10% neutral buffered formalin solution (Sigma-Aldrich, HT5012) for 1 minute. The reaction was quenched by washing with freshly prepared Tris-Glycine buffer twice, for 5 minutes each. Another PBS wash was performed, and tissue sections were incubated with 100mM NHS-Acetate mixture for 15 minutes at room temperature to quench reactive amine groups. After a 2X saline-sodium citrate (SSC) wash, the CosMx 6k Discovery Panel (Nanostring, 121500041) was applied. This pool of recently denatured and crash cooled in-situ hybridization probes targeting 6175 human genes plus 20 negative control probes was added along with RNase Inhibitor, Buffer R, and rRNA segmentation markers, and incubated overnight (18 hours) at 37°C in a RapidFISH Slide Hybridizer (Boekel Scientific, 240200). The following day, two stringent washes with 50% formamide were performed in a 37°C water bath for 25 minutes each, and slides were washed in 2X SSC twice. Tissue sections were incubated with a DAPI nuclear stain for 15 minutes, washed in PBS three times, and then incubated with neural-specific cell segmentation markers (GFAP and Histone) for 1 hour. Three more PBS washes were done, and samples were incubated with 100mM NHS-Acetate mixture for 15 minutes at room temperature again. Slides were washed with 2X SSC and flow cells were applied before CosMx machine loading. For the flow cell configuration setup, Configuration B was used for the pre-bleaching profile, and Configuration B was used for the cell segmentation profile, which was described as ideal for “human and mouse neural tissue”. Following the initial scan, approximately 1000 square fields of view (FOVs) were placed over the regions of interest and the run was initiated as described in the CosMx® SMI Instrument manual (Bruker Spatial Biology, MAN-10161-08).

**CosMx spatial transcriptomic data computational analysis**

Following scan completion and upload to NanoString's cloud AtoMx Spatial Analysis Platform, "studies" were created for each slide, and the following flat CSV files were exported for in-house analysis: count matrix, cell metadata, transcripts, polygons, and FOV positions. Scanpy (https://doi.org/10.1186/s13059-017-1382-0) was utilized for pre-processing, transcript-count-based filtering, normalizing, dimensionality reduction, and for visualizing the dataset. Cells were filtered by total transcript (probe) counts, excluding low-quality cells with <300 transcripts and high-count outliers with >7500 transcripts. For RNA expression visualization, data values were processed using MAGIC (https://doi.org/10.1016/j.cell.2018.05.061) to ensure uniformity of gene expression. Clusters were selected using gene expression thresholds. LGE representation was defined by ASCL1 enrichment over NKX2-1, with values filtered above 0.15 to remove false positives. MGE representation was defined by NKX2-1 enrichment over ASCL1, with values filtered above 0.15 to remove false positives. CGE representation was defined by NR2F2 expression above 0.15. Selected genes were then quantified within these clusters by flattening expression across all cells within the cluster and plotting boxplots, and spatial representation was performed using the 'sc.pl.embedding' function.

To enable consistent cell type annotation and to compare CosMx spatial transcriptomes and morphologies across developmental stages, CosMx datasets (24 GW and 35 GW) were integrated with a matched snm3C-seq reference (subsets of 2T and 3T) using a joint low-dimensional embedding. For integration, snm3C-seq gene body methylation signal was first converted into a pseudoexpression representation and inverted to match RNA directionality. For 35 GW preprocessing specifically, additional filtering was applied to remove a noticeable stripe of low-UMI cells prior to integration. Following filtering, data were normalized and log-transformed, and PCA (n_comps=50) was computed on the combined dataset. Harmony batch correction was performed on the PCA embedding using the dataset identity as the batch variable. A shared nearest-neighbor graph was then computed from the Harmony-corrected representation using n_pcs = 20 and n_neighbors = 15, followed by UMAP visualization. Cell type identities were assigned to CosMx cells by transferring snm3C-derived labels using nearest-neighbor mapping in the integrated embedding space, generating hierarchical transferred label sets used throughout downstream analyses. Transferred labels were retained only when the label transfer confidence score exceeded 0.6, with lower-confidence assignments flagged as uncertain (42% of cells). To validate transferred annotations, marker gene expression was visualized by cell type as z-scored pseudobulk heatmaps across selected populations. Spatial localization of major neuronal and glial subclasses and MSN subtypes was assessed by plotting cell centroids in tissue coordinate space with the target cell type highlighted and all other cells shown in grey for anatomical context. Finally, segmentation-derived nuclear area measurements present within the CosMx dataset were extracted per cell and compared across cell types and developmental stages by computing cell-type-level summary statistics and confidence intervals, enabling quantitative assessment of nuclear size differences between MSN subtypes, glial populations, and vascular-associated cell classes across developmental stages.

***Immunohistochemistry***

Tissue sections were cryosectioned at 30 μm, mounted onto Superfrost Plus slides, and stored at −80 °C until use. Frozen sections were baked overnight at 65 °C and subsequently rehydrated in TNT buffer (PBS containing 0.05% Triton X-100). Sections were post-fixed in 4% paraformaldehyde for 20 min, followed by incubation in 1% hydrogen peroxide in PBS for 30 min to quench endogenous peroxidase activity. Antigen retrieval was performed by heating slides in 10 mM sodium citrate buffer (pH 6.0) at 95–100 °C for 10 min, after which sections were allowed to cool at room temperature for 30 min. Sections were then incubated again in 1% hydrogen peroxide in PBS for 1 h and blocked for 1 h in Tris–NaCl blocking buffer (TNB; 0.1 M Tris–HCl, pH 7.5; 0.15 M NaCl; 0.5% blocking reagent, PerkinElmer). Primary antibodies diluted in TNB were applied overnight at 4 °C. Slides were subsequently incubated with either biotinylated secondary antibodies (1:250 in TNB) or Alexa Fluor–conjugated secondary antibodies for 2.5 h at room temperature. For signal amplification of biotinylated antibodies, horseradish peroxidase–conjugated streptavidin (1:200 in TNB) was applied for 30 min, followed by incubation with a tyramide-conjugated fluorophore (Cy5, Akoya), diluted 1:100 in amplification buffer (PerkinElmer), for 5 min. DAPI was included during secondary antibody incubation to label nuclei. Sections were washed with a TNT buffer between all steps. Following staining, sections were dehydrated, mounted with slide mounting medium, and coverslipped.

Representative high-resolution images of basal ganglia regions were acquired using a Stellaris laser scanning confocal microscope (Leica Microsystems) using the LASX software. Excitation was performed using a single line 405nm laser for the DAPI channel and a white light laser set at 495 nm, 590 nm, and 649 nm for the other channels, with laser power ranging from 1–10% across all channels. All images were acquired at a scan speed of 600 Hz in bidirectional mode and collected with HyD detectors with gain below 40. Images were captured using either a 20X 0.75NA Plan apochromat objective, with a final pixel size of 1.52 µm and a z-step of 0.04 µm, or a 40XOil 1.3NA Plan Apochromat objective, at a final pixel size of 0.48 µm and a z-step between 10-12.5 µm. Maximum intensity projections from z stacks are shown in the figures.

***MERFISH+ Gene Selection and Imaging:***

**MERFISH+ 463-gene probe library design and construction**

We developed custom primary probes using a Python-based workflow, following a previously established method[2] in which each gene was targeted by a set of 5 to 60 unique 40-mer sequences complementary to its mRNA. We then used transcript sequences from the human reference genome (hg38). Genes that were either too short or highly homologous to other targets were excluded, resulting in final panels of 463- genes. To each 40-mer, we appended three unique 20-nt readout sequences per gene, along with universal PCR priming sites at both ends (sequences see Table S1).

**Design of chromatin probes**

We designed probes for DNA hybridization similarly as those for the RNA MERFISH as described in Kern et al[2]. Briefly, we first partitioned *NKX2-1 and DRD2* locus into 10-kb segments. After screening against off-target binding, GC content and melting temperature,

~100 unique 40 bp target sequences were selected for each 10-kb segment. We concatenated a unique readout sequence to the target probes of each segment to facilitate sequential hybridization and imaging of each locus. Sequences are shown in Table S2.

#### Linker probes and codebook construction

Gene-specific linker probes comprised a 20-nt region complementary to each gene's readout sequence, followed by two tandem 20-nt amplification-binding sequences. Linkers were synthesized by IDT (sequences in Table S3). For MERFISH imaging, we assigned spatial barcodes using a binary codebook generated with a custom Python script (https://github.com/epigen-UCSD/MERFISH_Plus_Paper). Each gene received a randomly selected N-bit barcode containing exactly four "1" bits, while enforcing a minimum Hamming distance of four between barcodes to enable error correction. We then optimized the codebook with a Metropolis–Hastings algorithm to reduce co-occurrence of high-expression genes within the same hybridization round, guided by single-cell RNA-seq expression profiles[2].

#### Amplification and readout probes

Amplification probes were designed as a two-tier system. Level-1 probes carried a 20-nt arm complementary to the linker overhang and four binding sites for Level-2 probes. Level-2 probes hybridized to Level-1 and contained four additional sites for fluorescent readout probes. All amplification oligos were synthesized by IDT (sequences see Table S3). Readout probes were 20-nt oligos double-labeled with channel-specific dyes (Cy3/561 nm, Cy5/638 nm, Alexa Fluor 750/750 nm) and purchased from IDT with HPLC purification. (sequences see Table S3).

#### Acrydite probe synthesis

Acrydite probes were generated from oligonucleotide pools as previously described (doi: https://doi.org/10.1101/2025.11.02.686137). Briefly, Twist Biosciences oligopools were amplified by limited-cycle qPCR (~15–20 cycles) using 0.6 µM of each acrydite primer to produce DNA templates (sequences see Table S3). Templates were transcribed into RNA using T7 in vitro transcription (NEB, E2040S), then reverse-transcribed to single-stranded

DNA with 16 µM acrydite primers. RNA was removed by alkaline hydrolysis, and ssDNA oligos were column-purified (Zymo, D4060). Probes were stored at −20 °C.

**RNA MERFISH+ sample preparation**

RNA MERFISH+ samples were prepared as previously described[2]. Briefly, fresh-frozen human brains were cryosectioned at −20 °C on a Leica CM3050S. Serial 16 µm sections were collected and mounted on 40-mm round #1.5 coverslips pretreated with salinization and poly-L-lysine (Millipore, 2913997; Bioptechs, 0420-0323-2). Sections were fixed in 4% paraformaldehyde, at room temperature for 20 minutes, then permeabilized in 70% ethanol at 4C for 3 hours and then equilibrated in hybridization wash buffer (40% formamide in 2× SSC) for 10 min. Encoding probes were hybridized in 50 µL buffer (2× SSC, 50% formamide, 10% dextran sulfate) containing 5 µM encoding probes and 1 µM acrydite-modified dT+/LNA anchor probe (IDT) at 47 °C for 18–24 h. Samples were washed in 40% formamide/2× SSC/0.5% Tween-20 (30 min, room temperature), post-fixed in 4% paraformaldehyde (in 2× SSC), rinsed in 2× SSC with murine RNase inhibitor, and embedded in a thin 4% polyacrylamide gel to anchor RNAs[2]. MERFISH imaging was performed on a home-built system as described in Kern et al[2].

**Chromatin tracing sample preparation**

Sample preparation largely followed the RNA MERFISH+ tissue workflow, except that permeabilization was performed with detergent as described[1]. Briefly, fresh-frozen brains were cryosectioned and mounted on 40-mm round #1.5 coverslips pretreated with salinization and poly-L-lysine (Millipore, 2913997; Bioptechs, 0420-0323-2). Fixed sections were permeabilized with 0.5% Triton X-100 (Sigma-Aldrich, T8787) in 1× PBS supplemented with RNase inhibitors for 10 min at room temperature, then rinsed once in 1× PBS with RNase inhibitors. Coverslips were transferred to a fresh 60-mm Petri dish and treated with 0.1 M HCl (Thermo Scientific, 24308) for exactly 5 min, followed by three washes (5 min each) in 1× PBS with RNase inhibitors. Samples were equilibrated in hybridization wash buffer (50% formamide in 2× SSC) for 10 min at room temperature. Encoding probes were hybridized by applying 50 µL of hybridization buffer (50% formamide, 2× SSC, 10% dextran sulfate) containing 5 µM encoding probes and 1 µM acrydite-modified dT+/LNA anchor probe (IDT). Samples were denatured at 90 °C for exactly 3 min, then incubated in a humidified chamber at 47 °C for 18–24 h. After hybridization, sections were washed in 40% formamide/2× SSC/0.5% Tween-20 for 30 min, post-fixed in 4% paraformaldehyde in 2× SSC. Finally, samples were embedded in a 4% polyacrylamide gel as described for RNA MERFISH+.

**Imaging and adaptor/readout hybridization protocol**

MERFISH+ measurements were conducted on a custom microfluidics-microscope system with the configuration previously described (PMID: 38400890 ; **doi:** https://doi.org/10.1101/2025.11.02.686137 ), To enable multi-modal imaging we first sequentially hybridized fluorescent readout probes and then imaged the targeted genomic loci and then the targeted mRNAs. Specifically the following protocol was used in order:

1. 43 or 24 rounds of hybridization and chromatin tracing imaging, sequentially targeting the NKX2-1 using 1-color imaging; 24 rounds for 3-color DRD2 imaging.
2. 15 rounds of hybridization and MERFISH imaging, combinatorially targeting 463 genes using 2-color imaging

Following each hybridization the sample was imaged and then the signal was removed by flowing 80% formamide for 20 minutes and then re- equilibrating to 2XSSC for 10 minutes.

### Data Analysis

**RNA MERFISH+ image pre-processing: Flat-field correction:** To correct spatially non-uniform illumination, we computed a per-pixel median intensity map from the first imaging round for each fluorescence channel. Raw images were then normalized by inversely scaling each pixel by the corresponding median value. **Deconvolution:** Normalized images were deconvolved using Wiener filtering with a microscope-specific, custom point-spread function (PSF) to suppress noise and sharpen diffraction-limited signals. A high-pass filter (Gaussian blur $\sigma = 30$) was subsequently applied to further enhance spot contrast and improve signal-to-noise. **Spot localization:** Local intensity maxima within a 1-pixel neighborhood were detected and retained only if their brightness exceeded 3600 and their PSF correlation was ≥0.25. These filtered maxima were used for downstream decoding.

#### MERFISH decoding

To reconstruct RNA molecules across imaging rounds, fluorescent spots were first drift-corrected and then clustered across rounds by grouping spots located within 2 pixels of one another. Because the MERFISH codebook used a Hamming weight of 4, we retained clusters containing ≥3 spots to enable single–bit error correction, and discarded clusters with 1–2 spots. For each retained cluster, we assembled a per-round brightness vector, L2-normalized it, and matched it to the codebook using the same decoding strategy implemented in MERlin. To remove spurious calls, each decoded transcript was assigned a quality score derived from three features: (i) mean spot brightness, (ii) distance of the normalized brightness vector to the nearest codeword, and (iii) mean spatial dispersion of spots relative to the cluster median. For each feature, we computed a p-value relative to its global distribution across decoded transcripts and combined p-values using Fisher's method. Finally, we selected a score cutoff by comparing the score distributions of blank-barcode decodes versus gene-assigned decodes and choosing a threshold that separates their peaks.

#### Cell segmentation

Cell boundaries were segmented in 3D using Cellpose v2 (PMID: 36304142) or Cellpose V4. For each z-slice, we generated a 2D mask from the DAPI channel using the Cellpose “dapi” model after flat-field correction and deconvolution. Masks were then linked across adjacent z-slices to form 3D cell segmentations.

**Assigning transcripts to cells**

Transcripts detected by MERFISH were assigned to cells after correcting molecule coordinates for the measured drift between the segmentation images and the RNA imaging rounds. Corrected coordinates were rounded to the nearest pixel and mapped to the corresponding cell by querying the segmentation mask value at each transcript position.

**Chromatin tracing analysis**

**1) Localization of Fluorescent Spots**

To calculate fluorescent spot localizations for chromatin tracing data, we followed the following computational steps as described[2]: We first computed a microscope-specific point spread function (PSF) and, for each color channel, generated a per-pixel median illumination image from the first imaging round across all fields of view to normalize spatial intensity variation. For spot detection, images were flat-field corrected per channel, deconvolved using the custom PSF, and fluorescent puncta were identified as local maxima in the processed images.

**2) Image registration and selection of chromatin traces**

Image registration was achieved by aligning the DAPI channel for each field of view across imaging rounds. After flat-field correction and deconvolution, we identified local maxima and minima in the DAPI signal. A rigid translational offset was then estimated by FFT-based cross-correlation to optimally match these features between rounds. Nuclear segmentation was performed as described in RNA MERFISH+ above. Following image registration, chromatin traces were computed from the drift-corrected local maxima of each imaged locus as previously described (**doi:** https://doi.org/10.1101/2025.11.02.686137).

**Code availability**

Custom code used for analyzing chromatin tracing datasets in this study are available here: https://github.com/epigen-UCSD/MERFISH_Plus_Paper

**map3C Pipeline**

Snm3C-seq sequencing reads were trimmed, demultiplexed, and mapped to hg38 using the map3C pipeline[3].

**Quality Control and Preprocessing**

Cells were filtered on metadata thresholds; global CG methylation level > 0.5, global non-CG methylation level < 0.2, and total 3C interactions > 50,000. CG and CH methylation fractions were computed for 100-kb bins spanning the entire genomes as well as gene body methylation fractions where gene bodies were defined as annotated gene loci from gencode v33 flanked by 2kb on each end. Features with mean coverage less than 10 or intersecting with a methylation blacklist (https://github.com/Boyle-Lab/Blacklist/blob/master/lists/hg38-blacklist.v2.bed.gz) were removed. Additionally 10-kb CG methylation features were computed across the entire genome where each 10-kb bin is binarized as either methylated or not based on a methylation rate threshold of .9. Single-cell contact maps were first filtered using Hi-C 1D and 2D blacklists (https://github.com/zhoujt1994/scHiCluster/tree/master/files/blacklist). Contact matrices at 100 kb, 25kb and 10kb resolution were imputed by scHiCluster[4] with pad = 1,2 and 2 respectively. The imputed contacts with distance of >100 kb and <1 Mb were used as features for singular value decomposition dimension reduction. 3CGS is defined as the sum of off-diagonal contact values across rows and columns bounded by the transcription start site and transcription end site (TES) bins, with 2-kb flanking regions on each end, in the imputed 10-kb contact matrices.

**Embedding, Clustering, and Annotation**

A joint embedding of the 100kb 3C and 10-kb methylation features was created by concatenating the 100 10-kb PCs multiplied by 45, to create equal variances in each respective PC space, to the 50 3C PCs, then computing a neighbors graph with scanpy's[5] preprocessing neighbors function for 100 neighbors. A 2D UMAP was then built on this neighbor's graph. Cells were partitioned along unsupervised leiden clustering axes aligned the CG hypomethylation patterns of known canonical marker genes [6] to define L1 major type and L2 major lineages. Fine tuned L3 cell types were defined by reclustering the respective L2 major lineage partitions within each age group, and again by over batch correcting across ages using harmony for cell type consistency across ages.

**DMR**

All CG methylation DMRs were identified from pseudobulk allc files using Methylpy[7] (https://github.com/yupenghe/methylpy) with –num-sims=1000. Lineage DMRs were computed as the pairwise difference between any two adjacent developmental points, whereas branching DMRs were defined by pairwise comparisons between consecutive cell states along diverging developmental trajectories. daughter cell type to the parent cell type.

**Pseudotime**

Diffusion based pseudotime (DPT) analyses were computed using the methods outlined in ref. [8] Multi-modal lineage pseudotimes were computed using only cells from the newly generated data, excluding Tian 2023 and Heffel 2024, to mitigate pseudotime artifacts driven by batch effects. Each L2 lineage was reprocessed, computing PCA, k-nearest neighbors with k=50 using the top 50 principal components, for its respective subset of the data and limited to the Mid-gestation to 7 month age range to focus on development. The same cell was set as a root node in each modality and was selected from the earliest developmental cell type in each lineage as the cell furthest from the next adjacent cell type in the lineage. Genes were selected for display based on the highest positive and negative correlations to pseudotime as well as one vs all ranked t-test for each stage in a trajectory. Displayed genes were further limited to protein coding only.

**Compartment, Domain & Loop**

**Compartment**

Pseudo-bulk raw .cool files at 100-kb resolution were used for compartment analysis. We first constructed a reference contact map by merging contact maps from 5,500 randomly selected high-coverage cells and filtered out genomic bins with abnormal coverage. Following a procedure similar to that described in Tian et al[9], the merged contact map was used to fit a principal component analysis (PCA) model. The first principal component (PC1) was used as the bin-level compartment score, and its sign was adjusted such that compartments with higher CpG density had positive scores. For each age- and cell-type–specific pseudo-bulk matrix, contact maps of each major cell type were filtered and converted into correlation matrices in the same manner as described above, and were subsequently projected onto the fitted PCA model to obtain compartment scores.

Compartment strengths were computed as described in Tian et al[9]. Differential compartments were identified using dcHiC across major cell-type developmental lineages. Raw bin-level compartment scores were used to identify A–B and B–A compartment switches. We selected the top differential compartments based on a Z-score–transformed Mahalanobis distance greater than 1.96, corresponding to the 97.5th percentile of the standard normal distribution.

**Domain**

Chromatin domains were identified using single-cell imputed contact matrices at 25-kb resolution. Domain boundaries were defined at the single-cell level, and the boundary probability of each genomic bin was calculated as the proportion of cells in which the bin was called as a domain boundary relative to the total number of cells in the group. Insulation scores for each cell group were computed using pseudo-bulk imputed contact matrices.

To identify differential domain boundaries between pairwise comparisons, we constructed a 2×2 contingency table for each 25-kb bin based on the probabilities of cells being called as boundaries in each group. For each bin, we computed a chi-square statistic and corresponding P value, and defined differential domain boundaries as peaks of the chi-square statistic across the genome. Peaks were identified as local maxima of the chi-square statistic with a false discovery rate (FDR) < $1 \times 10^{-3}$ using the Benjamini–Hochberg procedure. In addition, candidate peaks were required to have a Z-score–transformed chi-square statistic greater than 1.960 (corresponding to the 97.5th percentile of the standard normal distribution), a fold change greater than 1.2 between the maximum and minimum insulation scores, and a difference greater than 0.05 between the maximum and minimum boundary probabilities.

**Loop**

Chromatin loops were identified through scHiCluster[4] pipelines following the same procedures as in our previous studies[1]. Loop calling was restricted to genomic distances between 50kb and 5Mb. For each single cell, imputed contact matrices were log-transformed and Z-score normalized along each diagonal to obtain globally normalized matrices (E_cell). Local background was estimated from interactions at distances of 30–50 kb and subtracted from E_cell to generate locally normalized matrices (T_cell). At the pseudo-bulk level, t statistics were computed across single cells to quantify deviations of E_cell and T_cell from zero. A null distribution was estimated by diagonal-wise shuffling of E_cell, from which T_shuffle and corresponding t statistics were derived, and empirical FDRs were obtained by comparing observed and shuffled statistics. Loop pixels were required to have an average E value > 0, fold changes >1.33 relative to donut and bottom-left backgrounds and >1.2 relative to horizontal and vertical backgrounds, and an empirical FDR < 0.01 against both global and local backgrounds. Loop summits were identified by clustering significant loop pixels within 20 kb using a breadth-first search algorithm and selecting the pixel with the highest E value in each cluster. Differential loops were identified between age groups in the same major lineage, following the approach described in the previous publications[1,9].

**ChromHMM analysis**

To characterize spatial and combinatorial patterns of methylation and/or chromatin contacts across various cell types and time points, we used the ChromHMM v1.26[10,11] software to conduct HMM state-based analysis. We trained two HMMs separately: the first model only included methylation features, and the second model jointly included methylation and chromatin contact features.

**Model learning**

We used 180 pseudobulk snm3C samples with an average coverage of greater than 50 cells. Each pseudobulk is of a specific cell-age combination. We divided the genome into bins and for each pseudobulk sample separately binarized the methylation and chromatin contact signals. For CpG methylation, a bin was encoded with a 2 corresponding to missing if it contained less than R reads, otherwise it was encoded as 1 if more than M% of its reads were methylated or 0 otherwise. For chromatin contact strengths, a bin was encoded with a 2 if the bin had low coverage or in the Hi-C blacklist regions, 1 if its signal was among the top 10% across all bins in the sample, or as 0 otherwise.  For the methylation only model, the bin size was 200bp, M=50 and R=3. For the joint model, the bin size was 5kbp, M=80 and R=75. We note that for the joint model we used a higher percentage threshold for binarization as combining 200bp bins into 5kbp reduced variance from the overall average methylation percentage of 84% causing few bins to have a methylation percentage of less than 50%.

We next trained these two types of HMMs using the "LearnModel" command of the ChromHMM[10,11] software using the gene-based binarized data files as input. For the methylation-only model, we used the "-b 200" flag to specify that the model's resolution is 200bp. For the joint model, we used the "-b 5000" flag to specify that the model's resolution is 5kbp. We trained both types of models with 5-80 states, with increments of 5. We focused our subsequent analysis on the largest models, which were the 80-state models.

**Overlap enrichment with universal ChromHMM states**

To investigate the overlap between the learned HMM states and the spatial-combinatorial patterns of chromatin marks across diverse tissue and cell types, we performed overlap enrichment analysis with the 100-state universal ChromHMM states[12]. We obtained the hg38 liftOver version of the annotations from Vu and Ernst[12] and used the "OverlapEnrichment" command of the ChromHMM software, with the "-b" flag to specify the models' resolutions respectively.

**Overlap enrichment with ChIP-Atlas**

To investigate the overlap between the learned HMM states and a large collection of cell-type-specific transcription factor (TF) binding profiles, we performed overlap enrichment analysis with the ChIP-seq experiments curated and uniformly processed as part of the ChIP-Atlas database[13]. We extracted 23,650 experiments of human TFs with the hg38 assembly that had at least 1000 peaks on the autosomal chromosomes, and for each experiment kept only those top 1000 peaks to make comparisons across experiments more uniform. We used the "OverlapEnrichment" command of the ChromHMM software, with the "-b" flag to specify the models' resolutions respectively and the "-center" flag to specify that the enrichments were computed based on the center base of each peak and the hypergeometric p-values were computed for the enrichments.

**DREM analysis**

To model the regulatory dynamics of DNA methylation across developmental lineages, we used the DREM (Dynamic Regulatory Events Miner) software (v2.0.6). Previous applications of DREM required two primary inputs: TF-gene interaction data and time-series expression data. We adapted the DREM framework to analyze methylation dynamics at differentially methylated regions (DMRs) rather than gene expression.

**TF-DMR interaction data generation**

We constructed TF-DMR interaction data using the ChIP-Atlas enrichment results from the 200bp resolution ChromHMM model described above. We selected the top 3 enriched ChIP-Atlas experiments per HMM state, yielding 176 unique experiments. For the target regions, we used 200-bp bins defined by merging DMRs called using the differential methylated sites (DMS) ≥ 2 criterion, which identifies regions exhibiting significant methylation changes across conditions. We intersected each ChIP-Atlas peak set with the DMR coordinate, retaining all experiment-DMR overlaps. Each ChIP-Atlas experiment was mapped to its corresponding TF and cell type using ChIP-Atlas metadata, and experiments were maintained as separate entities (keyed by TF, cell type, and accession number) rather than collapsed by TF identity. This preserved the cell type specificity of TF binding information in the final interaction matrix provided to DREM.

To enhance biological relevance to brain development, we filtered the TF list based on expression in the BrainSpan developmental transcriptome atlas. For each TF in our interaction data, we computed the mean RPKM across all BrainSpan samples and retained only those TFs with mean RPKM greater than a specified threshold of 1. This filtering step removed TFs with minimal brain expression that were less likely to have regulatory roles in the developmental context under study, while retaining TFs with consistent expression across brain development.

**DMR methylation dynamics quantification**

To generate the time series input for DREM, we computed DMR level methylation summaries at each developmental time point along a given lineage. For each time point, we extracted CpG level methylation data from the snm3C methylation profiles, recording the methylated cytosine count (mC) and total coverage (C_total) for each CpG site. We then aggregated CpG-level statistics within each DMR by summing mC and C_total values across constituent CpGs, and computed the methylation fraction (beta) as mC_sum/C_total_sum. We retained only DMRs with C_total_sum > 2 for more reliable beta estimates. The resulting beta matrix contained DMRs as rows and lineage time points as columns. Given that the 200bp resolution DMR set substantially exceeded the typical ~20,000 protein-coding genes for which DREM was designed, we filtered to the top 50,000 most dynamically changing DMRs, ranked by the range of beta values (maximum minus minimum) across the lineage.

**Model Parameters**

We applied DREM with a log transformation to input values, where it computes the log ratio relative to the first time point for each DMR. We set the minimum absolute log ratio threshold to 0.35 (compared to the default of 1.0) to capture the more subtle changes typical of methylation dynamics relative to gene expression. The penalized likelihood node penalty was set to 750 (compared to the default of 40) to account for the larger and highly dynamic input set of DMRs. TF associations at bifurcation points were assessed using path significance conditional on split, with a DREM score threshold of $10^{-4}$. The DREM score is computed with the hypergeometric distribution and is analogous to a p-value with smaller values reflecting a more confident association, but since the TF-DMR interaction was also used to learn a model and influence path assignments the score should not be interpreted as a p-value. All other parameters were set to default values.

**Met-scDRS to integrate GWAS with single cell methylomics**

Using met-scDRS [14] software and its released putative gene score, we applied the same 75 genome wide association studies (GWAS) spanning brain, immune/blood, metabolism, and other traits from previous publication [14,15] to single-cell methylome. The details on the derivation of gene score is detailed in the previous publication[14]. Briefly, these putative gene scores ($W_g$) are derived from GWAS summary statistics using MAGMA2 [16] with 10 kb windows flanking a gene body, summarizing the evidence of nearby genetic variants associated with the disease for that gene. Using recommended default parameters, methylation ratios ($X_{(c,g)}$) aggregated at the gene body were first inverted as ($[1 - X]_{(c,g)}$) and arcsine normalized. Genes with $< 5^{th}$ percentile in variance are filtered out for variance stabilization followed by regression of global methylation level as batch effect. Next, for each putative score ($W_g$), met-scDRS repeatedly sampled control genes with matching gene length, variance and normalized ratio and computed cell-wise disease association z-score (met-scDRS) and its p-value for each trait.

For each of the 75 traits, we then quantified the proportion of significant cells (FDR-corrected p-value < 0.1) in each L1, L2, and L3 cell types and visualized them using ComplexHeatmap3. Additionally, they are also further partitioned by developmental time points for further visualization in SCZ. For SCZ and ADHD, we visualized the met-scDRS with all available developmental time points as well as 1 month time point in UMAP latent dimension.

Finally, we prioritized genes with met-scDRS where genes with the 100 highest correlations between preprocessed methylation fraction and met-scDRS are identified between developmental time points and all time points for ADHD and SCZ respectively. We used these prioritized genes to perform gene ontology enrichment analysis using ClusterProfiler4 and visualized the genes memberships in enriched pathways.
